# MicroProteins miP1b/BBX30 and miP1a/BBX31 form a positive feedback loop with ABI5 to retard seedling establishment

**DOI:** 10.1101/2022.06.21.497044

**Authors:** Deeksha Singh, Sourav Datta

## Abstract

In plants the switch to autotrophic growth involves germination followed by post-germination seedling establishment. When environmental conditions are not favorable, the stress hormone Abscisic Acid (ABA) signals plants to postpone seedling establishment by inducing the expression of the transcription factor ABI5. The levels of ABI5 determine the efficiency of the ABA mediated post-germination developmental growth arrest. The molecular mechanisms regulating the stability and activity of ABI5 during the transition to light are less known. We found that two microProteins miP1a/ BBX31 and miP1b/BBX30 alongwith ABI5 inhibit post-germination seedling establishment in a partially interdependent manner. MicroProteins are single-domain proteins that interact with multi-domain proteins to modulate their post-translational activity. miP1a/ BBX31 and miP1b/BBX30 physically interact with ABI5 to stabilize it and promote its binding to promoters of downstream genes in light. ABI5 reciprocally induces the expression of *BBX30* and *BBX31* by directly binding to their promoter. ABI5 and the two microProteins thereby form a positive feedback loop to promote ABA-mediated developmental arrest of seedlings. Our study highlights the functional versatility of microProteins which can act as regulators of crucial developmental transitions in plants as well as other eukaryotes.

## Introduction

Growth is a fundamental characteristic of all living organisms. However, organisms invest energy in growth only after their survival is ensured. In seed plants, just after germination, the seed needs to make a crucial decision whether to proceed with seedling development or not, based on the environmental conditions. This decision involves an interplay of signals modulated by endogenous hormones and external factors. A post-germination developmental checkpoint arrests the growth of seedlings when conditions are not favorable. One of the key phytohormones that regulates germination and early plant development is Abscisic Acid (ABA)(Weitbrecht et al., 2011; Lopez-Molina et al., 2001). It is also known as the stress hormone that inhibits plant growth during times of stress. Among the external factors that regulate seed germination and post-germination development, light plays a crucial role(Xu et al., 2014). The role of ABA and light in regulating germination, indicated by the emergence of the radicle, is quite well studied. Seedling establishment, as determined by the presence of green, open and expanded cotyledons, has traditionally been studied as part of the germination process. However, several studies indicate that these two consecutive steps are developmentally distinct and regulated by common as well as unique factors(Lopez-Molina et al., 2001). Light promotes the autotrophic establishment of seedlings(Neff and Volkenburgh, 1994; Chattopadhyay et al., 1998; Deng et al., 1991; Chen et al., 2008a). The molecular understanding of the interplay between ABA and light to regulate post-germination seedling establishment is limited.

Abiotic stress signals induce the accumulation of ABA by elevating its biosynthesis and inhibiting its catabolism. ABA-deficient mutants germinate and establish faster than wildtype, whereas ABA catabolism mutants accumulate more ABA and exhibit longer dormancy periods (Tuteja, 2007; Xiong et al., 2002; Shu et al., 2018). ABA-induced by stress is perceived and channeled through a signaling pathway to induce changes in gene expression that ultimately lead to different stress responses. Studies utilizing the exogenous application of ABA have allowed us to understand better the ABA signaling pathway and ABA-induced responses. The ABA-signaling pathway involves the binding of ABA to PYRABACTIN RESISTANCE1 (PYR)/ PYR1-LIKE (PYL)/ REGULATORY COMPONENTS OF ABA RECEPTORS (RCAR) receptors leading to its interaction with type 2C protein phosphatases (PP2Cs). This interaction and retention of PP2Cs by ABA-bound receptors leads to the phosphorylation and release of the kinases SnRK2s, which further phosphorylate downstream transcription factors inducing ABA-mediated gene expression (Ali et al., 2020). Numerous transcription factors in Arabidopsis have been characterized for their roles in ABA signaling and response. The bZIP transcription factor ABSCISIC ACID INSESNITIVE5 (ABI5) was identified in a forward genetic screen for mutants exhibiting ABA insensitivity during germination (Finkelstein and Lynch, 2000). Although ABI5 regulates the ABA-mediated inhibition of germination, it plays a relatively more crucial role in desiccation tolerance and ABA sensitivity during post-germination development (Maia et al., 2014). ABI5 promotes post-germination arrest of seedlings to inhibit their growth and establishment under stress(Lopez-Molina et al., 2001). It binds to the G-box type ABA response elements (ABRE) on the promoters of several target genes including its promoter. EARLY METHIONINE-LABELLED 1 (EM1) and EM6, which code for LATE EMBRYOGENESIS ABUNDANT (LEA) proteins, are some of the key downstream targets of ABI5 (Choi et al., 2000; Finkelstein and Lynch, 2000; Carles et al., 2002).

Sensitivity to ABA during post-germination seedling development is substantially modulated by light(Yadukrishnan and Datta, 2021). The interactions between the ABA and the light signaling pathway components are reciprocal and multi-layered. ABI5 forms the point of convergence of interactions with light signaling factors like PHYTOCHROME-INTERACTING FACTORs (PIFs), ELONGATED HYPOCOTYL5 (HY5), FAR-RED ELONGATED HYPOCOTYLS3 (FHY3), DE-ETIOLATED 1 (DET1), and B-box (BBX) proteins at the transcriptional level (Yadukrishnan and Datta, 2021). In the dark, the PIF proteins – PIF1, PIF3, PIF4, and PIF5 – directly bind to the *ABI5* promoter and induce its transcription(Qi et al., 2020). Additionally, ABI5 physically interacts with PIF1 and enhances the binding of PIF1 onto the promoters of common target genes(Kim et al., 2016). The transcription factor HY5 also directly binds to the promoter of *ABI5* and promotes its expression(Chen et al., 2008a). It was recently identified that HY5 acts downstream of COP1 to negatively regulate ABA-mediated inhibition of postgermination seedling development(Yadukrishnan et al., 2020). FHY3 and DET1 directly bind to the *ABI5* promoter to activate or suppress its expression respectively and thereby optimize seedling greening during early development(Tang et al., 2013). In darkness, DET1 stabilizes the PIF proteins to indirectly enhance *ABI5* expression (Dong et al., 2014). BBX proteins are B-box containing zinc finger transcription factors that regulate several aspects of light-mediated development including photomorphogenesis, flowering, shade avoidance, high light, and UV-B tolerance (Gangappa and Botto, 2014; Xu et al., 2014). Some BBX proteins play crucial roles in the light-ABA signaling crosstalk(Vaishak et al., 2019). BBX21 is a transcriptional activator of *HY5* and a positive regulator of photomorphogenesis(Datta et al., 2007; Xu et al., 2014). BBX21 physically interacts with HY5 and inhibits its binding on the *ABI5* promoter(Xu et al., 2014). BBX21 also interacts with ABI5 and prevents it from binding on its own promoter, thereby reducing *ABI5* expression(Xu et al., 2014). Moreover, BBX21 directly binds to the *ABI5* promoter and recruits a chromatin remodeler protein HRB2 (HYPERSENSITIVE TO RED AND BLUE 2) to alter the chromatin structure on the *ABI5* promoter and reduce *ABI5* transcription(Kang et al., 2018). Another B-box protein BBX19 binds to the promoter of *ABI5* and induces its expression to promote ABA-mediated inhibition of germination and seedling development(Bai et al., 2019). CO and COL4/BBX5 also regulate ABA signaling, although the mechanistic details need further investigation (Min et al., 2015) ABI5 function is tightly regulated post-translationally by several factors. The activity of ABI5 is turned on by SnRK2s through phosphorylation, while it gets deactivated through dephosphorylation by PP2Cs(Nakashima et al., 2009). Furthermore, its activity is fine-tuned through modifications such as ubiquitination, sumoylation, and S-nitrosylation(Stone et al., 2006; Albertos et al., 2015; Miura and Hasegawa, 2009).The stability and activity of ABI5 protein are also modulated by components of the light signaling pathway like COP1. *cop1* shows ABA hyposensitivity during post-germination seedling development(Yadukrishnan et al., 2020). COP1 mediates ABA-induced accumulation of ABI5 by physically interacting with ABA-hypersensitive DCAF1 (ABD1) that targets ABI5 for degradation. COP1 ubiquitinates ABD1 to promote its degradation and thereby enhances ABI5 protein stability in dark(Peng et al., 2022). Additionally, COP1 promotes the binding of ABI5 to its target promoters to inhibit seedling growth (Yadukrishnan et al., 2020).·Regulation by microProteins has emerged as a relatively new mechanism of post-translational control of protein abundance and activity in various organisms (Rodrigues et al., 2021; Kruusvee et al., 2022; Kruusvee and Wenkel, 2022; Kushwaha et al., 2022). MicroProteins are small, single-domain proteins generally less than 140 amino acids long that share sequence homology with multi-domain proteins (Hong et al., 2005; Dolde et al., 2018). ABI5 inhibits seedling establishment under unfavorable conditions when a seedling tries to emerge into the light from the darkness underneath the soil. The efficacy of this post-germination developmental growth arrest depends on the levels of ABI5 (Lopez-Molina et al., 2003). How the stability of ABI5 is regulated, especially under light, and factors modulating its activity is relatively less understood.

Here we found that the two light inducible microroteins BBX31/miP1a and BBX30/miP1b (Wu et al., 2020) and ABI5 arrest post-germination seedling development in an interdependent manner. ABI5 directly binds to the promoter of *BBX30* and *BBX31* and induces their transcription. BBX30 and BBX31 physically interact with ABI5 and promote its stabilization. BBX30 and BBX31 also enhance ABI5-mediated gene expression by promoting the binding of ABI5 on its target promoters. Taken together our study suggests that ABI5 and the microProteins BBX30 and BBX31 arrest seedling growth by a positive feedback mechanism.

## Materials and methods

### Plant materials and growth conditions

In this study, the accession of *Arabidopsis thaliana* used is Columbia-0 (Col-0). The mutant and overexpressor lines *bbx30, bbx31, BBX30OE, BBX31OE, abi5-8, and ABI5OE* have been described previously(Graeff et al., 2016; Yadav et al., 2019; Yadukrishnan et al., 2020; Nambara et al., 1995). The double and triple mutants used in this study were generated by genetic crossing. The growth conditions and ABA treatments were similar to those described previously (Yadukrishnan et al., 2020). In short, the seeds were surface sterilized with sodium hypochlorite and stratified in water for 3 days. The seed was then sown on 0.5x MS (-)sucrose plates containing 1% agar and transferred to 16 h/8 h light/dark cycles of 80 μmol m^-2^sec^-1^of white light and 22°C in a Percival (CU-41L4) growth chamber for the desired number of days.

### Plasmid construction

To generate His-BBX30 and His-BBX31 constructs, the full-length coding sequences of *BBX30* and *BBX31* were cloned into the *BamHI-EcoRI* sites of the pET-28a vector and pGEX4T vector respectively. To generate GST-ABI5 construct the full-length *ABI5* was cloned in EcoRI-XhoI sites of the pGEX-4T-1. To generate YFP^c^-BBX30 and YFP^c^-BBX31 construct, the coding sequence of *BBX30* and *BBX31* were amplified, and the cloned in pDONR207. The BP product was then cloned in destination vector pCL113. Similarly to generate YFP·-ABI5, full-length ABI5 was cloned in pDONR207, and the BP product was cloned in pCL112. To create Yeast two-hybrid plasmid, the full-length coding region of *ABI5, BBX30* and *BBX31* were cloned in pGBKT7 (BD) and pGADT7 (AD) respectively. To study the domain interaction, we cloned different domains of ABI5 in the AD vector and full-length BBX31, N-terminal, and C-terminal of BBX31 in the BD vector. For the luciferase experiment, the reporter construct was generated by amplifying 1kb promoter of the *ABI5* gene followed by cloning at the KpnI-PstI sites of the pGreen II 0800-LUC vector(Lin et al., 2018). The effectors were generated by amplifying the coding sequence of the *BBX30, BBX31, and ABI5* gene, followed by its cloning in the pCAMBIA1300 vector using gateway cloning. All primers used to create the above-mentioned constructs are listed in Supplemental Table S1. The constructs were confirmed via sequencing before use.

### Quantification of germination and seedling establishment

Seeds of different genotypes were grown and harvested at the same time. The seeds were sterilized and stratified in water for 3 days. The seeds were then inoculated on 0.5x MS plates devoid of sucrose and supplemented with 0μM, 0.5μM, 1μM, 1.5μM of ABA. For estimating % germination, the seeds with completely emerged radicle were counted whereas for % seedling establishment seedlings with open green cotyledons were considered. All germination and seedling establishment experiments were performed thrice with >100 seeds per experiment. All the observations were made and representative images captured using Leica S6E stereomicroscope (Leica Microsystems, www.leica-microsystems.com).

### Yeast two-hybrid assays

The protein interaction study was performed as described previously (Ravindran et al., 2021). Briefly, the respective combinations of BBX30, BBX31, and ABI5 were co-transformed into the yeast strain *AH109*. Yeast transformants were then grown on the selection medium lacking leucine and tryptophan (DDO/-Trp-Leu). The interaction was further confirmed by growing on a medium lacking Trp, Leu, Ade, and His (QDO/-Trp-Leu-Ade-His).

### *In-Vitro* pull-down assay

The His-BBX30, His-BBX31, and GST-ABI5 constructs were transformed into ArcticExpress (DE3) cells. For GST-ABI5, the secondary culture of ArcticExpress DE3 transformed cells was induced by 0.5mM IPTG at 28° C for 4 hours. The lysate was incubated with GST beads (Glutathione Sepharose 4B, GE Healthcare) for 2 hours at 4° C. In the case of His-BBX30 and His-BBX31, the cell lysate was incubated with Ni-NTA beads (Ni Sepharose 6 Fast flow, GE Healthcare) for 45 mins. The GST and HIS protein-bound beads were washed with 1XPBS+300mM NaCl and 1XPBS+300mM NaCl+20mM imidazole respectively two times. GST-ABI5 was eluted using 30mM glutathione and His-tagged proteins were eluted using 250mM imidazole. The eluted proteins were concentrated, and excessive glutathione and imidazole were removed using MERCK concentrators. The purified 0.5μM GST-ABI5 and GST were allowed to bind to the GST beads for 2h at 4° C. The protein-bound beads were then incubated with purified His-BBX30 and His-BBX31 for 2 hours at 4° C. The beads were then washed with PBS twice to remove the unbound protein. For other combination 0.5 μM of His-BBX30 and His-BBX31 were bound to the beads and used as bait, while prey proteins GST, and GST-ABI5 were allowed to bind to the bead bound bait proteins. The protein bound to the beads were separated using 10% SDS gel, followed by blotting using Anti-His (SAB4301134, Sigma) and Anti-GST antibody (ab9085, abcam).

### Bi-molecular fluorescence complementation assay

BiFC was performed as previously mentioned (Gampala et al., 2007). Briefly, the N-terminal and C-Terminal half of YFP vectors carrying full-length *ABI5, BBX30*, and *BBX31* respectively were transformed into Agrobacterium cells. The bacteria were grown overnight, briefly pelleted and resuspended in infiltration buffer (10mM MES, pH 5.7, 10mM MgCl2, 150mM acetosyringone). The suspensions for the constructs as required for the transformation were mixed and infiltrated into young, fully expanded leaves of *N. benthamiana*. The infiltrated leaves were imaged after 2 days using an FV-3000 Olympus confocal microscope with excitation of 488 nm and emission between 510-525 nm.

### Protein Co-localisation

Destination vector pGWB661 was used to generate BBX30-RFP and BBX31-RFP, while ABI5 was cloned in PGWB6 to generate GFP-ABI5. Three weeks old N. benthamiana were infiltrated with Agrobacterium cells transformed with BBX30-RFP and GFP-ABI5 or BBX31-RFP and GFP-ABI5. After 2days incubation in dark, the tagged proteins were visualised using an olympus live cell microscope. The excitation wavelengths of 488 nm and 561 nm was used for GFP and RFP, respectively.

### Immunoblotting

For Anti-ABI5 immunoblots, the germinated seeds of Col-0, *bbx30bbx31, BBX31OE, ABI5OE*, and *bbx30bbx31ABI5OE* were transferred to media containing 10 μM ABA for indicated time intervals and the harvested samples were crushed in liquid nitrogen and homogenized in extraction buffer (50 mM Tris–HCl, pH 7.5, 75 mM NaCl, 10 mM EDTA, 10 mM MgCl2, 0.1% Tween 20, 1mM NaF, and 1 X proteases inhibitor mix, Sigma) followed by centrifugation for 10 min at 11,000 g at 4 °C. Protein concentrations were determined by the Bradford method. 40μg of total protein were loaded per well in SDS-acrylamide/bisacrylamide gel and proteins were electrophoretically transferred to a PVDF membrane (Millipore). For treatment with MG132 and ABA, the germinated seeds of Col-0 and *bbx30bbx31* were transferred to media containing 0.5x MS -sucrose with 10μm ABA or 50μm MG132 for the indicated time. Samples were harvested and blotted using an anti-ABI5 antibody. Actin was used as an internal loading sample control. For cycloheximide (CHX) treatment, germinated seeds were treated with 20μM ABA for 12 hours in the presence of 100μM CHX and samples were harvested at different time intervals. H3 protein was used as a sample loading control. Proteins were detected using Anti-ABI5 and Anti-H3 antibody. For immunoblotting, the membrane was blocked for 1 hour in Tris-buffered saline-0.1% Tween 20 (TBST) containing 5% BSA. After blocking the proteins were probed with antibodies diluted in TBST overnight at 4° C. The antibodies used were Anti-ABI5 (ab98831; Abcam, www.abcam.com), Anti-ACTIN (Sigma A0480), Anti-H3 (AS10 710, Agrisera). The membrane was then washed thrice with TBST followed by secondary antibody incubation for 1 hour at room temperature. The secondary antibody used was goat-raised horseradish peroxidase-conjugated antirabbit IgG (Sigma) in 1:8000 dilution. To develop the membrane, ECL western blotting substrate (PierceTM), was used. The protein bands were visualized using the ChemiDoc MP imaging system (Bio-Rad). The band intensity was measured using ImageJ software.

### Electrophoretic mobility shift assay (EMSA)

Full-length *BBX30* and *BBX31* cloned in pCold-TF vector was transformed into ArcticExpress DE3 cells. The secondary culture was induced with 0.2mM IPTG at 28° C for 3 hours, and the cell lysate was incubated with Ni-NTA beads for 1 hour at 4° C. EMSA was performed as described previously(Job et al., 2018). Briefly, 30-50 base pair oligos were labeled using (Biotin 3’
s End DNA Labeling Kit; Pierce), followed by annealing of complementary strands. 1μg of mentioned proteins and 2μg in case of 2X, were incubated with 20 fmol biotinylated probes. The binding was checked by running the incubation mixture on 6% (v/v) native polyacrylamide gels in 0.5× TBE. The DNA-protein complex was blotted to a positively charged nylon membrane and imaged using Chemiluminescent Nucleic Acid Detection Module (Thermo Fisher Scientific). The primers used are listed in Supplemental Table S1.

### Chromatin immunoprecipitation (ChIP)-qPCR

The ChIP assay was performed as described previously (Saleh et al., 2008; Komar et al., 2016; Yadukrishnan et al., 2020). Briefly, approximately 1g of seeds were sterilized and grown on a 0.5xMS-suc plate containing 1μM ABA for 3 days. The harvested tissue was crosslinked by fixing in 1% formaldehyde along with vacuum infiltration. The crosslinking was stopped by adding 0.125M glycine to it. Next, chromatin was isolated using a series of extraction buffers as mentioned in (Saleh et al., 2008), followed by sonication using 4 cycles-30 sec on/30 sec off at 4° C. 10 percent of the total chromatin sample was stored as input, the rest chromatin sample was incubated with antibody-coated beads at 4° C. The bead-bound chromatin was then processed for reverse crosslinking using 5M NaCl and 65° C incubation for 4-6 hours. The beads were then removed from the solution using the magnetic rack, and the DNA was purified using a DNA fragment purification kit. The enrichment was checked through qPCR using primers specific to the promoter/protein binding region. The percentage input methods were used to analyze the enrichment. All the primers used are listed in Supplemental Table S1.

### Luciferase assay

The protoplast isolation was done as described earlier (Sheen, 2002). pGreen II 0800-LUC vector was used to generate the *ABI5·: LUC* reporter. *BBX30, BBX31*, and *ABI5* cloned under *35S* promoter were used as effectors. The vectors in different combinations were transfected into the protoplast via PEG solution and incubated overnight to ensure the expression of protein and the activation of the promoter. The luciferase activity was measured using the Promega kit (E1910) and *Renilla luciferase* activity was used as an internal control.

### RNA isolation and qPCR

RNA was isolated from germinated seeds as previously mentioned (Yadukrishnan et al., 2020). cDNA was prepared using Bio-Rad iSCRIPT cDNA synthesis kit as instructed in the manufacturer’s protocol. qPCR was performed in LightCycler®96 (Roche,www.roche.com) machine, and TB Green® Premix EX Taq·II SYBR Green dye was used (TaKaRa, www.takarabio.com). For internal control, the reference genes used were UBIQUITIN10 and GAPDH. The qPCR primers used are listed in Supplemental Table S1. The values depicted in the graphs represent the three independent experiments.

### Statistical analysis

All the statistical analyses were performed using Graph Pad Prism 9.0 and Microsoft Excel. To determine the statistical significance, one-way ANOVA was performed followed by Tukey’s *post hoc* test. Other details of the analyses are mentioned in the Fig. legends.

### Accession numbers

*BBX30* (AT4G15248), *BBX31* (AT3G21890), *ABI5* (AT2G36270), *EM1* (AT3G51810), *EM6* (AT2G40170).

## Results

### BBX30 and BBX31 promote ABA-mediated inhibition of post-germination seedling development

*BBX30* and *BBX31* are well-characterized light-responsive genes playing roles in photomorphogenesis, UV-B signaling, and flowering (Heng et al., 2019; Yadav et al., 2019; Graeff et al., 2016). In one of our previous studies, we found that the expression of several genes involved in ABA response and desiccation tolerance was upregulated in *35S:BBX31* (Yadav et al., 2019). In another study, similar genes were downregulated in *bbx30bbx31*(Heng et al., 2019). *BBBX30* and *BBX31* expression is also upregulated by ABA(Xu et al., 2019). The promoter sequences of *BBX30* and *BBX31* contain Abscisic Acid (ABA) response elements (ABRE). All of these prompted us to ask the question if these genes play a role in ABA-regulated developmental responses. We started studying early seedling development, considering the broad function of ABA in regulating germination and post-germination events. We grew imbibed seeds of Col-0, *bbx30, bbx31, bbx30bbx31, BBX30OE*, and *BBX31OE* on a half-strength MS plate, in the presence (+) or absence (-) of ABA. Seedlings with open, expanded green cotyledons were used to calculate % seedling establishment (Fig. S1a-b). In (-) ABA conditions we did not find significant difference in the % seedling establishment of Col-0, *bbx30, bbx31, bbx30bbx31, BBX30OE*, and *BBX31OE*. Seedlings of all the genotypes showed 100% seedling establishment by 3·day after stratification (DAS) (Fig. 1a-b,d). However, the % seedling establishment in +ABA conditions was higher in *bbx30, bbx31* and *bbx30bbx31* and substantially lower in *BBX30OE* and *BBX31OE* compared to Col-0 (Fig. 1a, c and e). In order to verify if the difference in seedling establishment rate is due to differences in germination, we determind the % germination by counting seeds with emerged radicle in (-) and (+) ABA conditions. However we did not find difference in % germination among the genotypes in either (-) or (+) ABA containing plates. Seedlings of all the genotypes exhibit 100% germination by the 2·day after stratification (DAS) in -ABA conditions while total germination is achieved at 4 DAS under +ABA conditions (Fig. 1f-i). To further confirm the role of BBX30 and BBX31 in ABA-mediated inhibition of seedling development post germination, we germinated seeds on -ABA plates and then transferred them to +ABA plates to determine seedling establishment rate. About 40% of Col-0 seedlings were able to establish in +ABA conditions (Fig. S2a-c). *bbx30, bbx31, bbx30bbx31* seedlings showed higher seedling establishment compared to Col-0, while the seedling establishment % was only 20% for *BBX30OE* and *BBX31OE* seedlings (Fig. S2a-c). This suggests that BBX30 and BBX31 promote ABA-mediated inhibition of seedling development at the post-germination stage.

**Fig. 1.**
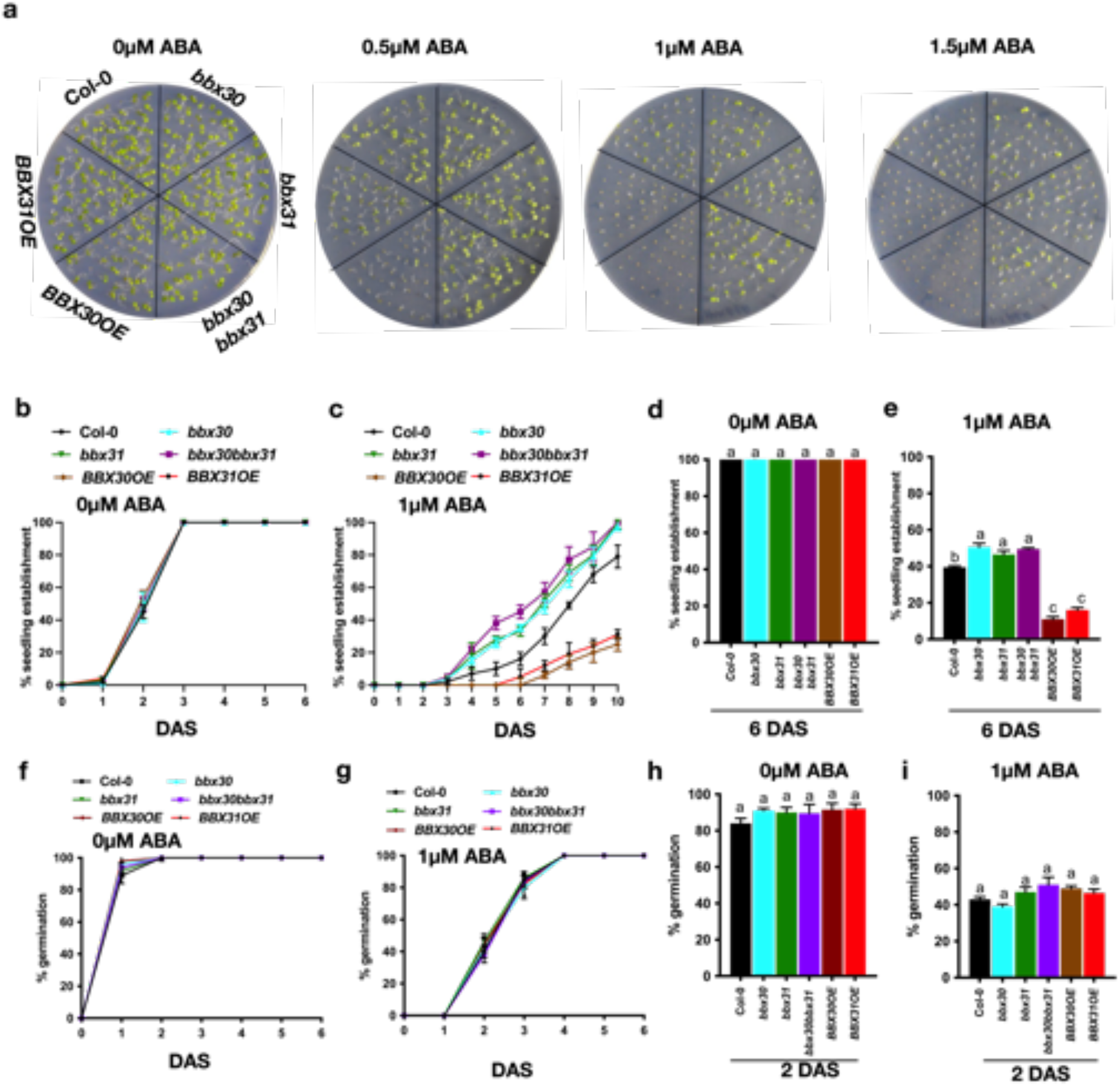
BBX30 and BBX31 promote ABA-mediated inhibition of post-germination seedling development. (a) Representative images of 6-day-old seedlings of Col-0, *bbx30, bbx31, bbx30bbx31, BBX30OE* and *BBX31OE* grown on a 0.5x MS plate supplemented with 0μM, 0.5μM, 1μM, 1.5μM of ABA. (b-c) % seedling establishment in (b) 0μM ABA and (c) 1μM ABA conditions for the indicated genotypes for up to 10 days after stratification (DAS) and on day 6 (d-e). (f-g) % germination of the indicated genotypes under 0μM (f) and 1μm (g) ABA conditions for up to 6 days after stratification (DAS) and on day 2 (h-i). Seedlings with green, open and expanded cotyledons were counted to determine % seedling establishment. In (b-i) Error bars represent SEM of three independent experiments with >500 seeds. In (d,e,h,i) the letters above the bar indicate the statistical groups as determined by one-way ANOVA followed by Tukey’s *post hoc* test (p < 0.05).

### BBX30, BBX31 and ABI5 mediate post-germination seedling growth arrest in an inter dependent manner

The transcription factor ABI5 is a key regulator of several ABA responses including post-germination development and is known to mediate interplay with light signaling factors (Chen et al., 2008b; Yadukrishnan et al., 2020; Lopez-Molina et al., 2001). We asked the question if ABI5 plays any role in BBX30 and BBX31 mediated seedling growth arrest. In the presence of ABA, the percentage of seedling establishment is higher in *abi5-8* compared to Col-0, *bbx30* and *bbx31* (Fig. 2a, c and e). This difference was not seen in -ABA conditions (Fig. 2a, b and d). In order to examine the genetic interaction between *BBX30, BBX31* and *ABI5* we generated *bbx30abi5-8* and *bbx31abi5-8* by crossing and monitored the seedling establishment rates of the double mutants under ABA. The establishment rates in *bbx30abi5-8*, and *bbx31abi5-8* were similar to *abi5-8* suggesting that BBX30 and BBX31 regulate these processes in an ABI5-dependent fashion (Fig. 2a-e). There was no difference in % germination among these genotypes in (-) ABA conditions, with all genotypes attaining 100% germination by 2 DAS (Fig. 2f and h). In (+) ABA conditions at 2 DAS % germination was higher in *abi5-8, bbx30abi5-8* and *bbx31abi5-8* compared to Col-0, *bbx30* and *bbx31*, however by 5 DAS there was no significant difference in the germination % between the genotypes (Fig. 2g and i).

**Fig. 2.**
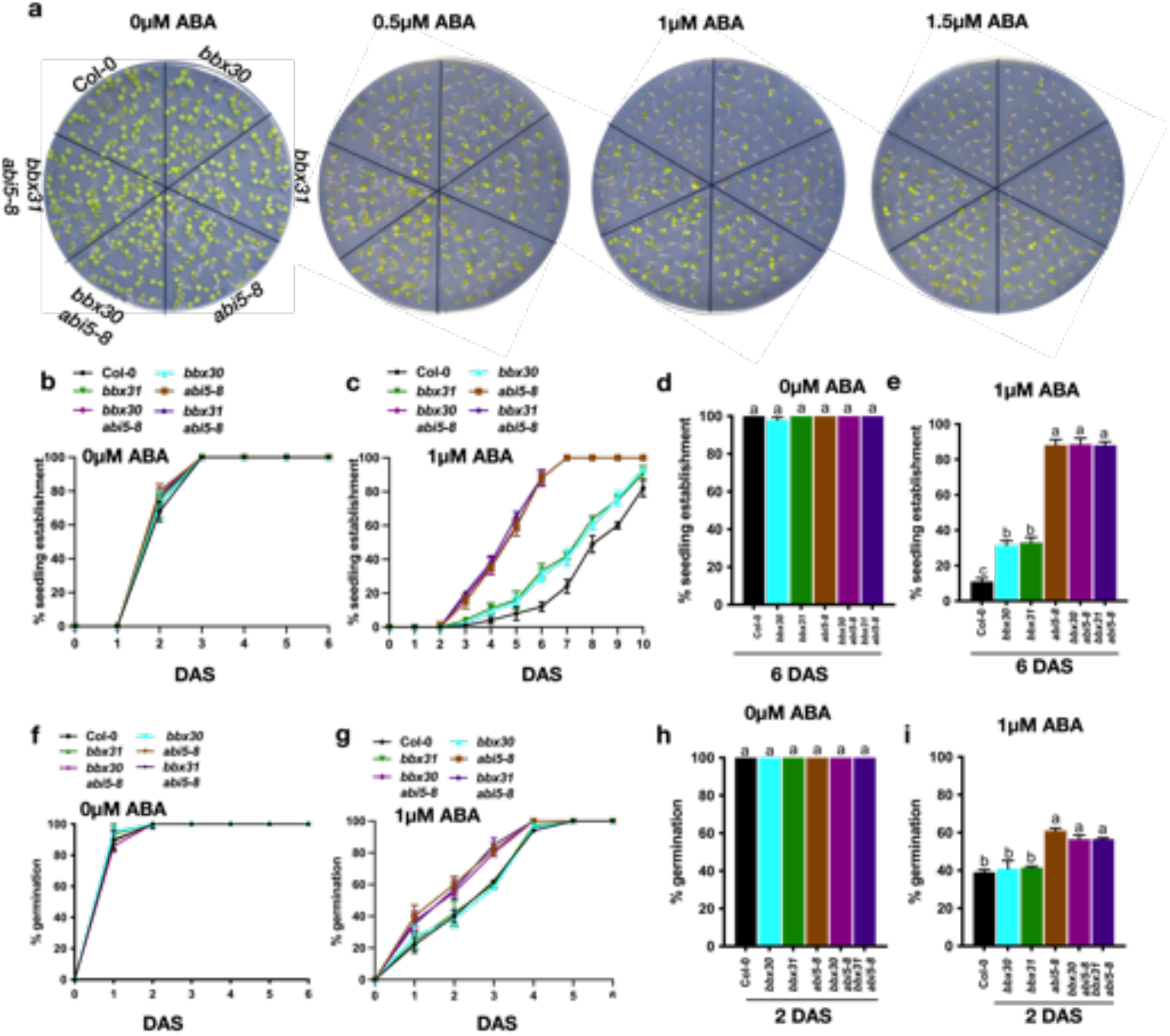
*BBX30, BBX31* and *ABI5* interact genetically to regulate early development in Arabidopsis. (a) Representative images of 6 day-old seedlings of Col-0, *bbx30, bbx31, abi5-8, abi5-8bbx30, abi5-8bbx31* grown on 0.5x MS plate supplemented with 0μM, 0.5μM, 1μM, 1.5μM of ABA. (b-c) % seedling establishment in (b) 0μM ABA and (c) 1μM ABA conditions for the indicated genotypes for up to 10 days after stratification (DAS) and on day 6 (d-e). (f-g) % germination of the indicated genotypes under 0μM (f) and 1μM (g) ABA conditions for upto 6 days after stratification (DAS) and on day 2 (h-i). Seedlings with green, open and expanded cotyledons were counted to determine % seedling establishment. In (b-i) Error bars represent SEM of three independet experiments with > 500 seeds. In (d,e,h,i) the letters above the bar indicate the statistical groups as determined by one-way ANOVA followed by Tukey’s *post hoc* test (p < 0.05).

We further crossed *BBX31OE* with *abi5-8* to obtain *abi5-8BBX31OE* and studied their ABA sensitivity during early seedling development (Fig. 3a). At 6 DAS under 1μM ABA, *abi5-8* and *BBX31OE* exhibit 85% and 2% seedling establishment respectively as compared to 15% in Col-0 (Fig. 3c-e). Seedling establishment % in *abi5-8BBX31OE* is around 40% (Fig. 3, c and e). There is no difference in the % seedling establishment between the genotypes under 0μM ABA (Fig. 3b-d). Germination rates of Col-0, *abi5-8, BBX31OE* and *abi5-8BBX31OE* by 5 DAS in presence or absence of ABA also did not show any significant difference (Fig. 3f-i).These data further validate that BBX31-mediated inhibition of post-germination seedling development is at least partially ABI5 dependent.

**Fig. 3.**
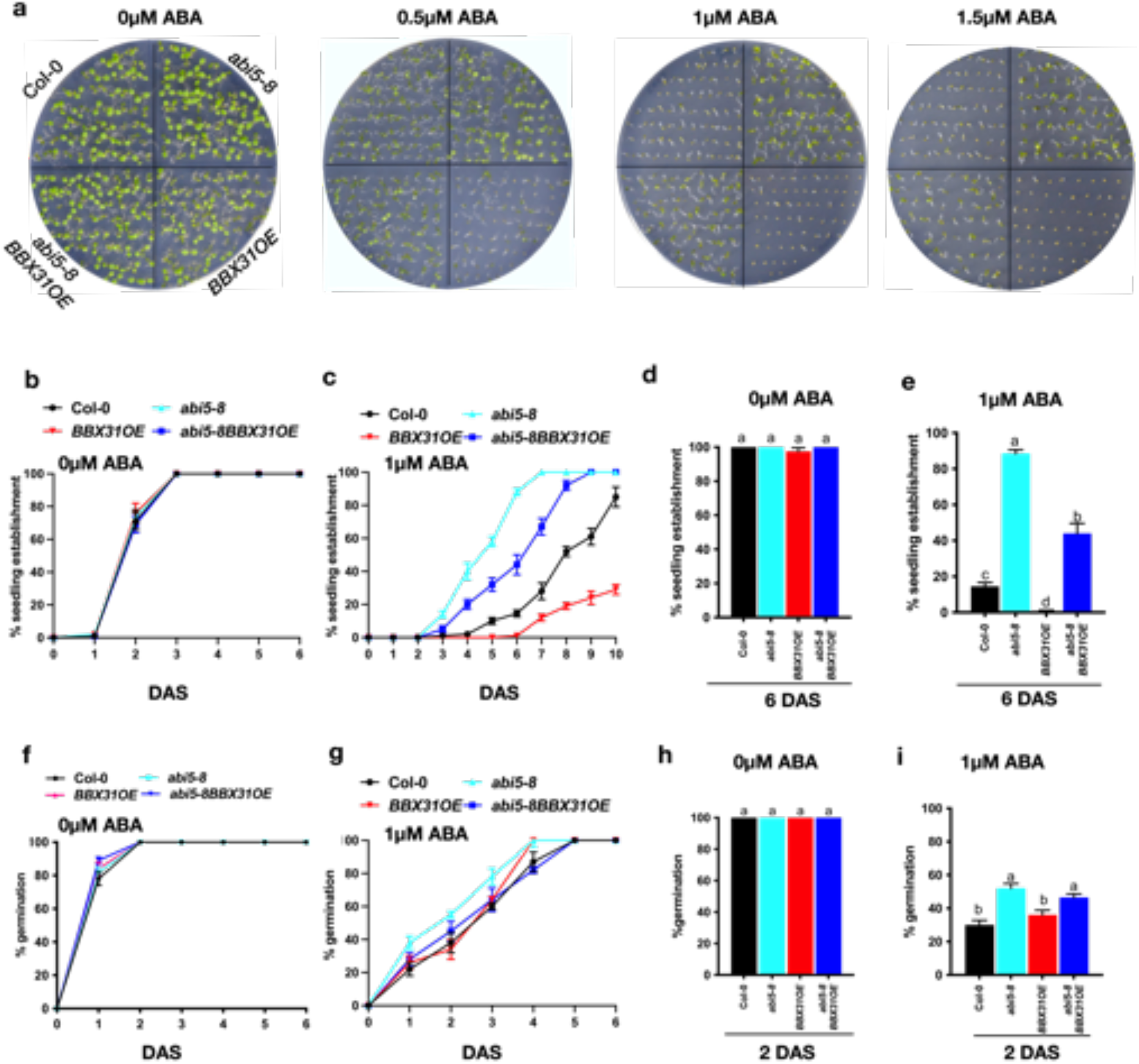
BBX31 regulates seedling growth arrest in an ABI5-dependent fashion. (a) Representative images of 6-day-old seedlings of Col-0, *abi5-8, BBX31OE, abi5-8BBX31OE* grown on a 0.5x MS plate supplemented with 0μM, 0.5μM, 1μM, 1.5μM of ABA. (b-c) % seedling establishment in (b) 0μM ABA and (c) 1μM ABA conditions for the indicated genotypes for up to 10 days after stratification (DAS) and on day 6 (d-e). (f-g) % germination of the indicated genotypes under 0μM (f) and 1μM (g) ABA conditions for upto 6 days after stratification (DAS) and on day 2 (h-i). Seedlings with green, open and expanded cotyledons were counted to determine % seedling establishment. In (b-i) Error bars represent SEM of three independent experiments with >500 seeds. In (d,e,h,i) the letters above the bar indicate the statistical groups as determined by one-way ANOVA followed by Tukey’s *post hoc* test (p < 0.05).

Next, we set up genetic crosses of *ABI5* overexpression line (*ABI5OE*) with *bbx30, bbx31*, and *bbx30bbx31* to generate *bbx30ABI5OE, bbx31ABI5OE*, and *bbx30bbx31ABI5OE* lines, respectively. Under 0μM ABA, germination and seedling establishment of all the genotypes progressed at similar rates till attaining 100% (Fig. 4a, b and C, Fig. S3 a and b). In presence of 1μM ABA, *ABI5OE* exhibited lower rate of seedling establishment as compared to the wildtype (Fig. 4a, d and e). Interestingly, *bbx30ABI5OE, bbx31ABI5OE*, and *bbx30bbx31ABI5OE* lines showed higher seedling establishment compared to *ABI5OE* (Fig. 4a, d and e). We observed similar growth pattern among the genotypes in 0.5 and 1.5 μm ABA (Fig. 4a). In 1μM ABA, *ABI5OE* showed lower % germination compared to Col-0, which was partially complemented in *bbx30ABI5OE, bbx31ABI5OE*, and *bbx30bbx31ABI5OE* lines at 2 DAS (Fig. S3 c, e). All these findings together suggest that the BBX proteins BBX30 and BBX31 are required for ABI5-mediated seedling growth arrest. This further suggests that the dependence of ABI5 on BBX30 and BBX31 is probably at the protein level, as the reduced seedling establishment phenotype of seedlings constitutively expressing b *ABI5* is partially rescued by the loss of *BBX30* and *BBX31*.

**Fig. 4.**
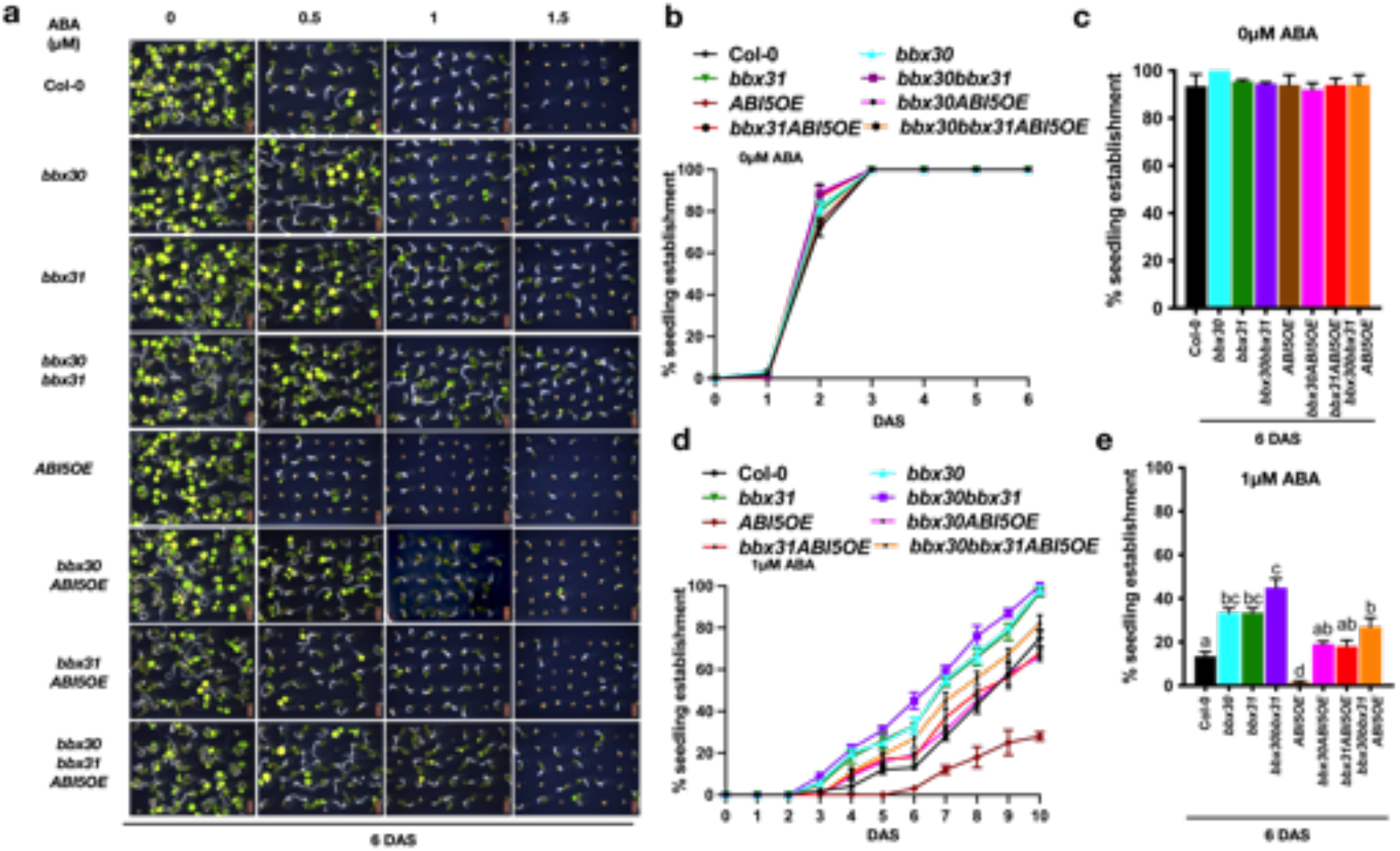
BBX30 and BBX31 are required for ABI5-mediated seedling growth arrest. (a) Representative images of 6-day-old seedlings of Col-0, *bbx30, bbx31, bbx30bbx31, ABI5OE, bbx30ABI5OE, bbx31ABI5OE, bbx30bbx31ABI5OE* grown on a 0.5x MS plate supplemented with 0μM, 0.5μM, 1μM, 1.5μM of ABA. (b-d) % seedling establishment in (b) 0μM and (d) 1μM ABA conditions for the indicated genotypes for up to 10 days after stratification (DAS) and on day 6 (c,e). Seedlings with green, open and expanded cotyledons were counted to determine % seedling establishment. In (b-e) Error bars represent SEM of three independent experiments with 500 seeds. In (c) and (e) letters above the bar indicate the statistical groups as determined by one-way ANOVA followed by Tukey’s *post hoc* test (p < 0.05).

### ABI5 physically interacts with BBX30 and BBX31

Since constitutive overexpression of *ABI5* could not enhance ABA response in the absence of *BBX30* and *BBX31*, we hypothesized that BBX30 and BBX31 might interact with ABI5 protein and regulate its activity. In order to confirm physical interaction between ABI5 and the BBX proteins we used yeast two-hybrid, in vitro pull down and BiFC assays. Our yeast-two-hybrid assay using ABI5 as bait and either BBX30 or BBX31 as prey indicated that ABI5 can physically interact with both BBX30 and BBX31 (Fig. 5a). According to previous studies ABI5 contains conserved domains, that govern specific functions (Lopez-Molina et al., 2003; Finkelstein and Lynch, 2000). We examined the interaction of BBX31 with different domains of ABI5, and observed that the conserved C1 domain of ABI5 is required for the interaction with the N-terminal of BBX31 (Fig. S4a and b). To further confirm the interaction between BBX30, BBX311 and ABI5, we performed in-vitro pull-down experiment using full length ABI5 fused to GST, and BBX30 and BBX31 proteins fused to 6xHis epitope. His-tagged BBX30 and BBX31 was able to pull down GST-ABI5, validating that BBX30 and BBX31 physically interact with ABI5 in vitro (Fig. 5b). Furthermore, we confimed the colocalisation of the two proteins using GFP tagged ABI5 and RFP tagged BBX30 and BBX31 coexpressed in *Nicotiana benthamiana* (Fig. S4c-d). We also validated the interaction of these proteins in vivo using bimolecular fluorescence complementation (BiFC) in *Nicotiana benthamiana* (Fig. 5c). All these data together suggest that ABI5 physically interacts with BBX30 and BBX31.

**Fig. 5.**
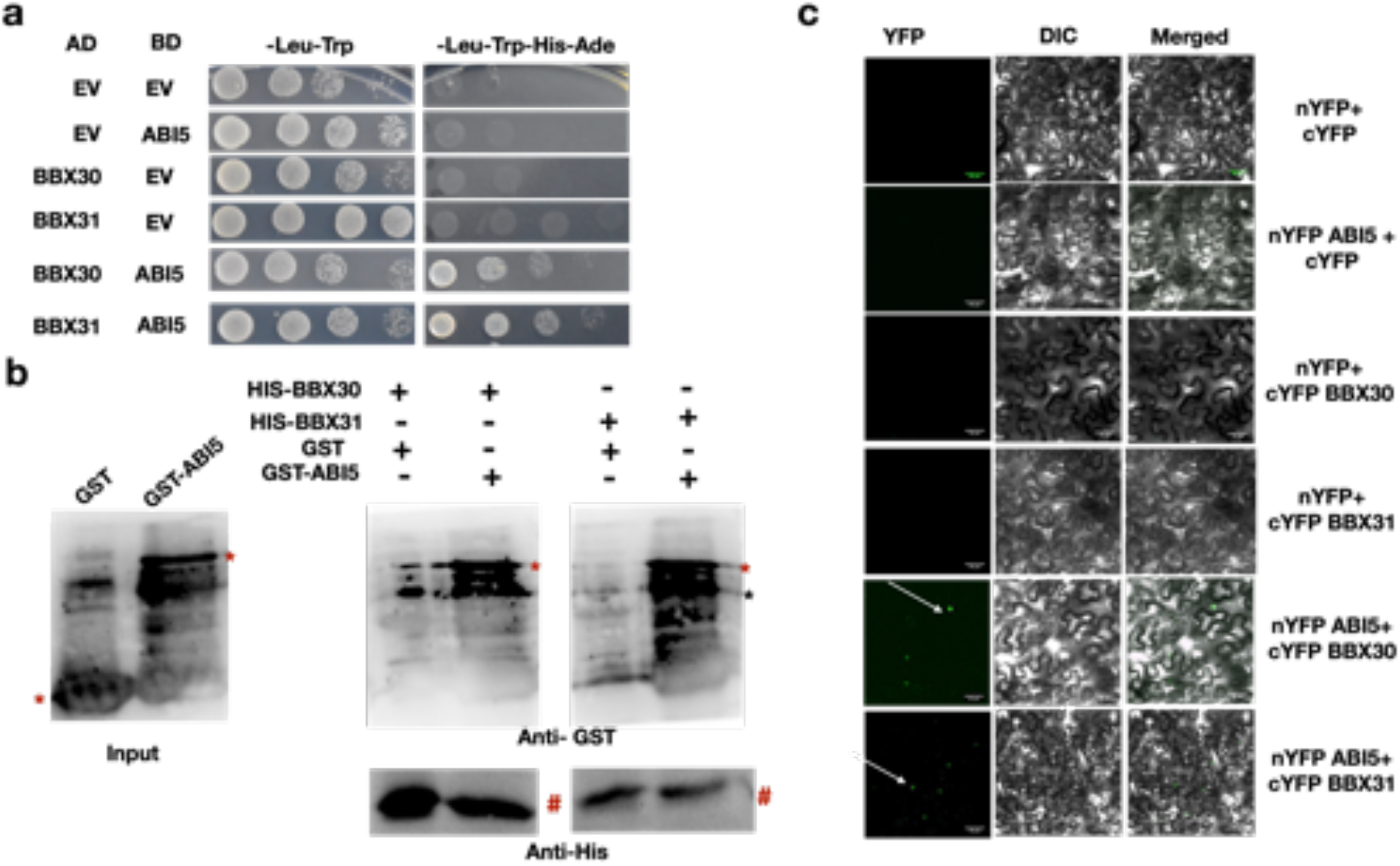
ABI5 physically interacts with BBX30 and BBX31. (a) Yeast two-hybrid assay showing the interaction between BBX30, BBX31 and ABI5. pGADT7 (AD) and pGBKT7 (BD) represent the *GAL4* activation domain and binding domain respectively. EV indicates empty AD or BD vectors. BBX30 and BBX31 were fused to AD, while ABI5 was fused with BD and tested for interaction. DDO represents the medium lacking leucine and tryptophan, while QDO additionally lacks histidine and adenine. (b) In-vitro pull-down assay showing the interaction between BBX30, BBX31 with ABI5. BBX30 and BBX31 were fused with N-terminal His tag, while ABI5 is fused with N-terminal GST tag. To check the physical interaction bead-bound His-BBX30 and His-BBX31 were used to pull down GST and ABI5-GST. Precipitated proteins were analysed by anti-GST and anti-His antibodies. *(red) mark indicate band specific to GST-ABI5 and # indicates Anti-His corresponding band, * (black) indicates nonspecific band. (c) BiFC assay showing the interaction of BBX30 and BBX31 with ABI5 in planta. nYFP and cYFP represent the N-terminal and C-terminal half of yellow fluorescent protein. Vectors in the indicated combinations were transformed into Agrobacterium and co-infiltrated into the 3-weeks-old *N. benthamiana* leaves and imaged after 36 hours. The scale bar represents 50μm.

### BBX30 and BBX31 stabilize ABI5 during post-germination development

BBX30 and BBX31 belong to the group of microProteins that can sequester other proteins and often regulate their stability (Wu et al., 2020; Graeff et al., 2016). Our protein-protein interaction studies indicated that BBX30 and BBX31 physically interact with ABI5 (Fig. 5). To understand the physiological significance of this interaction, we examined whether BBX30 and BBX31 can regulate the abundance of the ABI5 protein. Initially we put Col-0 and *bbx30bbx31* seeds on plates containing 0μM and 1μM ABA and harvested samples every day for up to 4 days after stratification and determind ABI5 accumulation by immunoblotting (Fig. S5a,b). ABI5 accumulated to high levels on day 1 in both Col-0 and *bbx30bbx31* (Fig. S5a,b). In -ABA conditions ABI5 accumulation in day 2, 3 and 4 samples in both the genotypes was decreased (Fig. S5a,b). In presence of ABA, ABI5 accumulation was detected on day 2, 3 and 4 samples in Col-0 but not in *bbx30bbx31* (Fig. S5a,b). This suggests that BBX30 and BBX31 might regulate the stability of ABI5 especially during post-germination development. To check the dynamics of ABI5 protein accumulation upon ABA treatment during post-germination development, we treated germinated seeds of Col-0, *bbx30bbx31*, and *BBX31OE* with 10μM ABA for 1, 6, and 12 hours and harvested samples to determine ABI5 protein levels (Fig. 6a,d). ABI5 accumulation was lower in *bbx30bbx31* compared to Col-0, whereas in *BBX31OE* enhanced accumulation of ABI5 was detected (Fig. 6a-d). Similar ABA treatment causes a gradual increase in the ABI5 accumulation in *ABI5OE*, while this is compromised in *bbx30 bbx31 ABI5OE*, suggesting that these BBX proteins promote ABA mediated post-germination ABI5 accumulation (Fig. 6e). Previous reports have shown that ABI5 protein undergoes degradation via the 26S proteasomal pathway (Lee et al., 2010; Stone et al., 2006). Inhibition of the 26S proteasome pathway by treating germinated seeds with MG132 for 6 hours prevented the decrease in ABI5 levels in *bbx30bbx31*, suggesting that BBX30 and BBX31 could promote the accumulation of ABI5 by negatively regulating its degradation (Fig. 6f). To uncouple the effects of de novo translation on ABI5 levels in these genotypes, we treated the germinated seeds with the translational inhibitor cycloheximide (CHX) and examined the stabilization of ABI5 in Col-0, *bbx30bbx31 and BBX31OE* (Fig. 6g and h). The rate of degradation of ABI5 was faster in *bbx30bbx31*, while reduced in *BBX31OE* as compared to Col-0 (Fig. 6g and h). All these results together suggest that BBX30 and BBX31 promote the stabilization of ABI5 during post-germination development by negatively regulating its proteasome-mediated degradation.

**Fig. 6.**
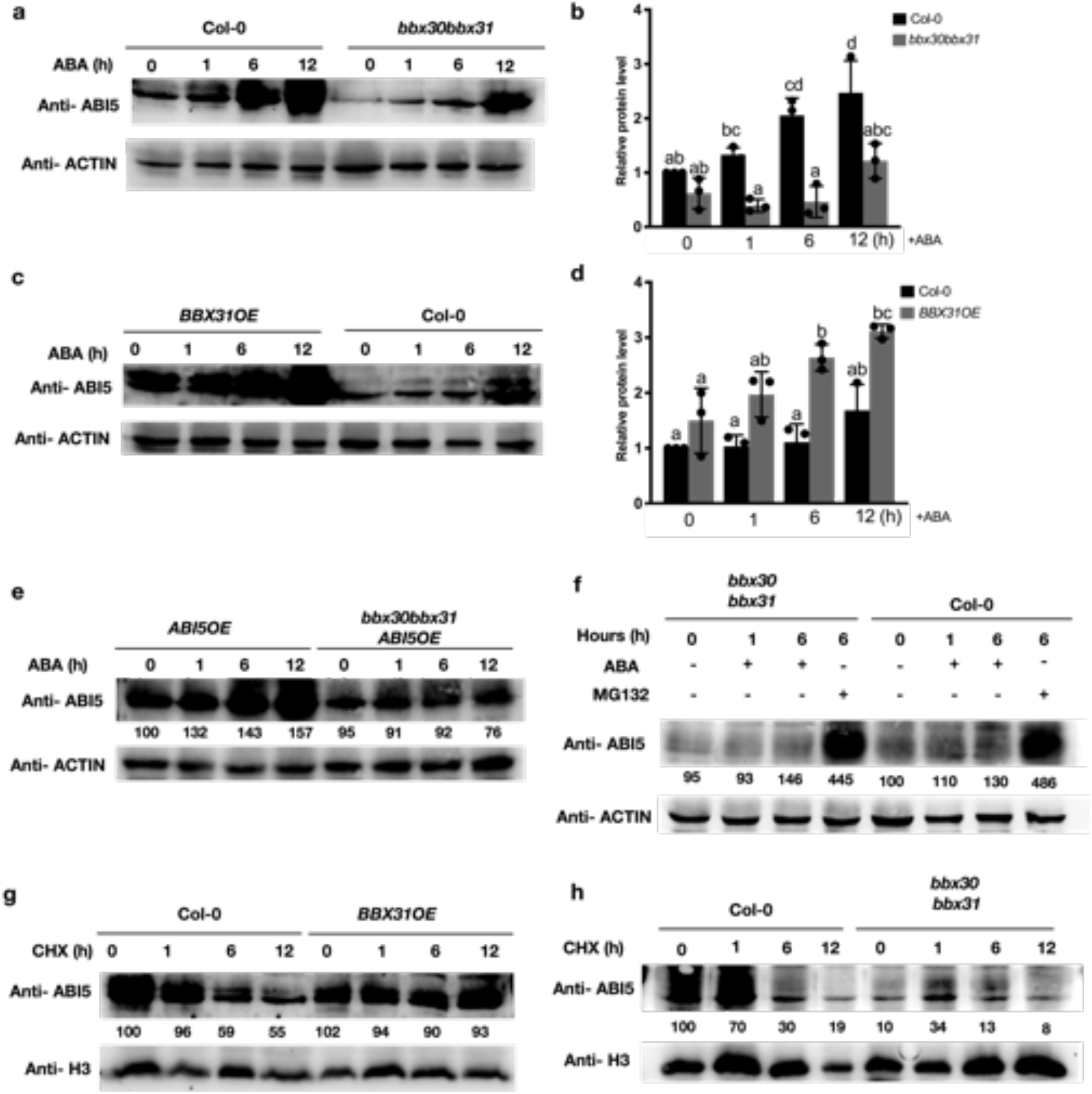
BBX30 and BBX31 stabilize ABI5. (a) Immunoblot showing the abundance of ABI5 protein in Col-0 and *bbx30bbx31*. (b) Relative protein levels of ABI5 in Col-0 and *bbx30bbx31*. (c) Immunoblot showing the abundance of of ABI5 protein in *BBX31OE* and Col-0. (d) Relative protein levels of ABI5 in *BBX31OE* and Col-0. In (b) and (d) error bars represent standard deviation of two blots. Letters above the error bar indicate the statistical groups as determined by one-way ANOVA followed by Tukey’s *post hoc* test (p < 0.05). (e) Immunoblot showing the protein level of ABI5 in *ABI5OE* and *bbx30bbx31ABI5OE* upon ABA treatment. In (a), (c) and (e) 1-day-old, germinated seeds were transferred from 0.5x MS -sucrose medium containing plates to similar plates with 10μM ABA for indicated time intervals and then harvested and subjected to immunoblotting using an anti-ABI5 antibody. Actin was used as an internal sample control and blotted using an anti-ACTIN antibody. (f) Immunoblot showing the level of ABI5 in *bbx30bbx31* and Col-0. Germinated seeds as in (a) were transferred to media containing 0.5x MS -sucrose with 10μm ABA or 50μm MG132 for the indicated time. Samples were harvested and blotted using an anti-ABI5 antibody. Actin was used as an internal loading sample control. (g-h) Immunoblot showing the protein levels of ABI5 in Col-0, *BBX31OE*, and *bbx30bbx31* upon CHX treatment. Germinated seeds were treated with 20μM ABA for 12 hours in the presence of 100μM CHX and samples were harvested at different time intervals. H3 protein was used as a sample loading control. Proteins were detected using Anti-ABI5 and Anti-H3 antibody respectively. In (e-h) numbers below each blot indicate the relative band intensity of proteins that is normalized to the loading control (ACTIN/H3). The intensity of the first band was set to 100.

### BBX30 and BBX31 enhance ABI5-mediated gene expression by promoting the binding of ABI5 on its target promoters

We now asked the question if BBX30 and BBX31 can modulate ABI5 activity. ABI5 is known to directly activate its own transcription and also a range of other target genes such as *EM1* and *EM6* (Chen et al., 2008c; Xu et al., 2014; Finkelstein and Lynch, 2000; Leung and Giraudat, 1998). First, we compared the expression level of *ABI5* and *EM6*, in Col-0, *bbx30bbx31, bbx30, bbx31, BBX30OE, BBX31OE* under -ABA and +ABA conditions in tissues of two different developmental stages - in germinated seeds that have completed the radicle emergence and in developing seedlings that have started but not completed the cotyledon emergence. We observed no significant difference in the expression of *ABI5* and *EM6* between the genotypes in the germinated seeds (Fig. 7a and c). However, the emerging seedlings of *BBX30OE* and *BBX31OE* showed higher expression of *ABI5* and *EM6* as compared to Col-0 in the presence of ABA, whereas the expression was lower in the loss of function mutants (Fig. 7b and d). Furthermore, we performed a transient assay in *Arabidopsis* protoplasts using luciferase gene driven by *ABI5* promoter as the reporter and ABI5, BBX30, and BBX31 as effectors (Fig. 7e). While the three effectors could individually induce the expression of *ABI5pro:LUC*, coexpression of *ABI5, BBX30*, and *BBX31* significantly increased the luciferase activity, suggesting that BBX30 and BBX31 enhance the transcriptional activation of target genes by ABI5 (Fig. 7f).

**Fig. 7.**
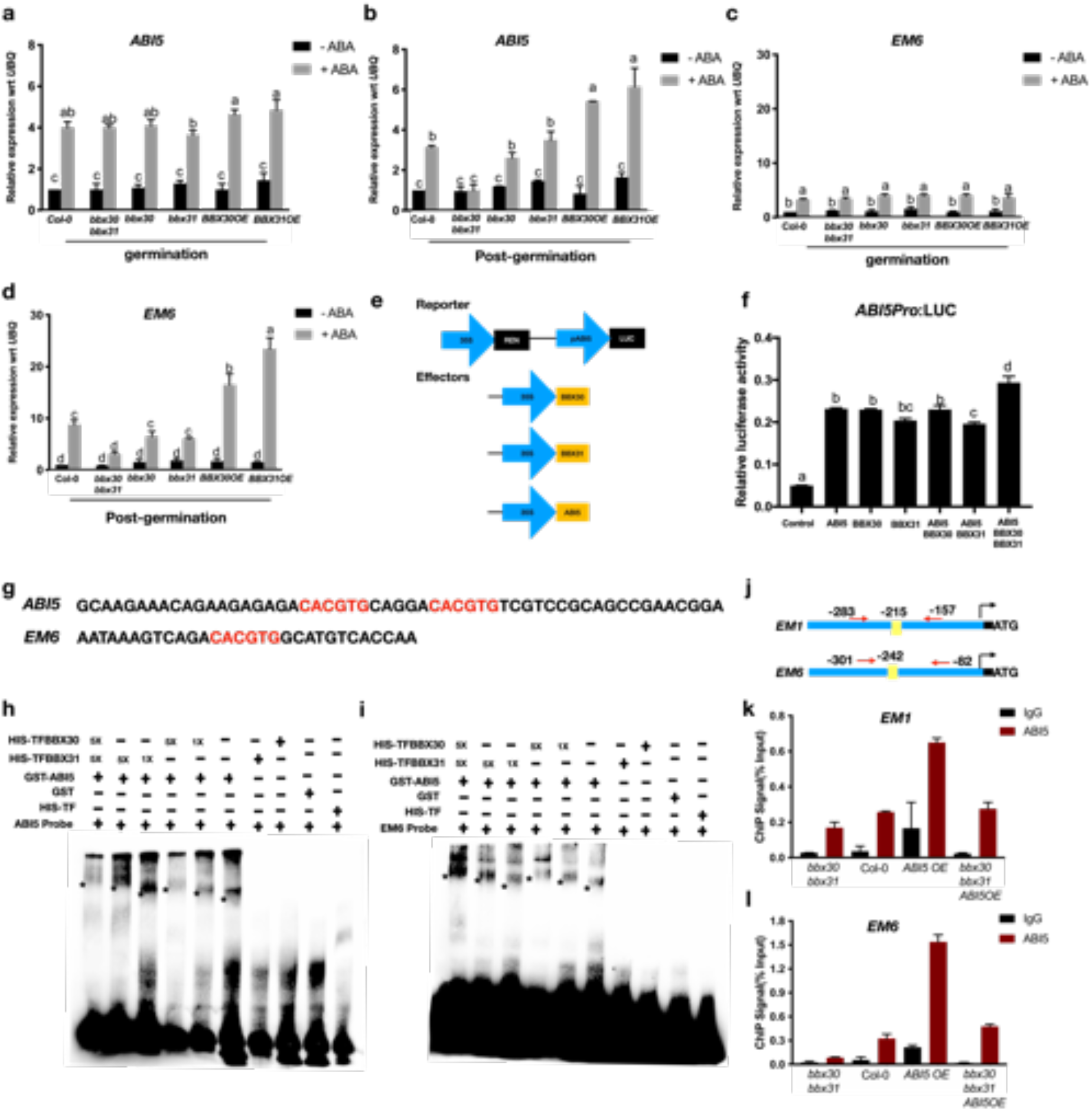
BBX30 and BBX31 enhance ABI5-mediated gene expression by promoting the binding of ABI5 on its target promoters. (a-d) Relative expression of (a-b) *ABI5* and (c-d) *EM6* in Col-0, *bbx30 bbx31, bbx30, bbx31, BBX30 OE, BBX31 OE*. Seeds were inoculated on 0.5x MS -sucrose plates, with or without 1μm ABA. The seeds were harvested on the second day (germination) and the third day (post-germination) after stratification. Values are the mean of three biological replicates. (e) Schematic representation of the constructs used in the transient assay in *Arabidopsis* protoplast. The reporter construct used is based on the pGreen vector expressing *35S promoter* driven Renilla Luciferase and *ABI5* _*pro*_ driven Luciferase. Effectors used are *ABI5, BBX30, BBX31* cloned under the constitutive *35S* promoter. (f) Relative luciferase activity, showing the activation of *ABI5*_*pro*_: LUC by the combination of proteins mentioned. Error bar represents SD (n=3). (g) The sequence of promoter regions of *ABI5* and *EM6* used as probes for the EMSA. The red highlighted region is the G-box type ABRE *cis*-regulatory element. (h-i) EMSA showing the binding of ABI5 along with BBX30 and BBX31. BBX30 and BBX31 alone, do not bind to *ABI5* and *EM6* promoters. The addition of BBX30 and BBX31 in the presence of ABI5 leads to a super shift (denoted by *) of the DNA-protein complex. + and - denotes the presence and absence of the probe or proteins indicated on the left. 1X and 5X represent the concentration of BBX30 and BBX31. (j) Schematic representation of *EM1* and *EM6* promoter. Black arrow represents the transcription start site. The yellow boxes indicates G-box type ABRE *cis*-regulatory elements, and the number represents the base location of G-box type elements. Red arrows denote primer binding sites for the primers used to check enrichment by qPCR following ChIP. (k-l) ChIP-qPCR analysis determining the binding of ABI5 onto the promoter of (k) *EM1* and (l) *EM6* in seedlings of *bbx30bbx31*, Col-0, *ABI5OE* and *bbx30 bbx31ABI5OE*. The genotypes used were grown in 1um ABA and samples were harvested on the fourth day after stratification. DNA-Protein complexes were immunoprecipitated using anti-ABI5 and anti-IgG antibodies (negative control). ChIP DNA was quantified using primers flanking the G-box region of *EM1* and *EM6*. Letters above the error bar (a-d and f) indicated the statistical groups as determined by one-way ANOVA followed by Tukey’s *post hoc* test (p < 0.05).

Subsequently, we performed an electrophoretic mobility shift assay (EMSA) to test the effect of BBX30 and BBX31 on the binding of ABI5 on its target promoters. BBX30 and BBX31 did not show direct binding on the promoter regions of *ABI5* and *EM6* where ABI5 is known to bind (Fig. 7g-i) However, the addition of BBX30 and BBX31 along with ABI5 resulted in a super shift as compared to ABI5 alone, indicating that BBX30 and BBX31 might form a complex with ABI5 during the latter’s DNA binding (Fig. 7h and i). To test whether BBX30 and BBX31 influence the binding of ABI5 on its target promoters in vivo, we performed a chromatin immunoprecipitation assay (ChIP) using anti-ABI5 antibody, followed by qPCR using primers flanking the ABI5-binding sites of *EM1* and *EM6* promoters in 3 DAS seedlings of Col-0, *bbx30 bbx31, ABI5OE*, and *bbx30 bbx31 ABI5OE*. The enrichment of ABI5 protein on *EM1* and *EM6* was decreased and increased in *bbx30bbx31* and *ABI5OE*, respectively, as compared to Col-0 (Fig. 7j-l). ChIP-qPCR showed the lack of ABI5 enrichment over *ACTIN* promoter used as a control (Fig. S6a). Interestingly, *bbx30bbx31ABI5OE* exhibited less ABI5 protein enrichment on *EM1* and *EM6* as compared to *ABI5OE*, suggesting that binding of ABI5 on the target promoters is promoted by BBX30 and BBX31 (Fig. 7j-l).

### ABI5 directly binds to the promoters of *BBX30* and *BBX31* and induces their transcription

*BBX30* and *BBX31* are well characterized as light-responsive genes induced during dark to light transition and are also regulated by UV-B, and the circadian clock (Yadav et al., 2019; Wu et al., 2020; Graeff et al., 2016). Since our data suggest the important role of BBX30 and BBX31 in ABA response during post-germination development, we asked whether these genes are transcriptionally regulated by ABA. We examined the expression levels of *BBX30* and *BBX31* by RT-qPCR in germinated Col-0 seeds treated with 10μM ABA for 0, 1, 6 and 12 hours (Fig. 8a-b). ABA treatment for 1 hour results in 3-fold and 12-fold upregulation in the mRNA levels of *BBX30* and *BBX31* respectively (Fig. 8a-b). Treatment with ABA for longer durations up to 12 hours results in the expression levels of *BBX30* and *BBX31* elevating to 4-folds and 15-folds compared to no ABA treatment (0 hour) (Fig. 8a-b). The G-box type ABA response elements (ABRE) present in the promoter of *BBX30* and *BBX31* are known to be the target sites of the transcription factor ABI5 that plays crucial roles in ABA signaling. To test if *BBX30* and *BBX31* are induced by ABA in an ABI5-dependent manner, we examined their expression in *abi5-8* treated similarly as explained above for Col-0. We found that the absence of ABI5 impairs the ABA-induced upregulation of *BBX30* and *BBX31* (Fig. 8a-b). All these data suggest that after germination ABA induces the expression of *BBX30* and *BBX31* in a ABI5 dependent manner. Next, we asked the question if ABI5 can directly bind to the promoter of *BBX30* and *BBX31*. To examine this, we performed EMSA using GST-ABI5 protein and biotinylated *BBX30* and *BBX31* promoter regions as the probes. GST-ABI5 was able to bind to the G-box-containing promoter sequences of *BBX30* and *BBX31*, while the binding was abrogated in the probes containing mutated G-boxes, indicating that ABI5 directly binds on the G-box elements on the promoters of *BBX30* and *BBX31* (Fig. 8c-e). To validate the in vivo binding of ABI5 on these promoters, we also performed a ChIP-qPCR, and examined the regions of *BBX30* and *BBX31* promoters showing ABI5 enrichment. The results indicated that ABI5 indeed binds to *BBX30* and *BBX31* promoters on the regions containing the G-box type ABRE cis-regulatory elements (Fig. 8f-h). These evidences suggest that while BBX30 and BBX31 interact with ABI5 to stabilize it and promote the expression of downstream ABA responsive genes, ABI5 establishes a positive feedback loop to directly activate the transcription of *BBX30* and *BBX31* (Fig. 9).

**Fig. 8.**
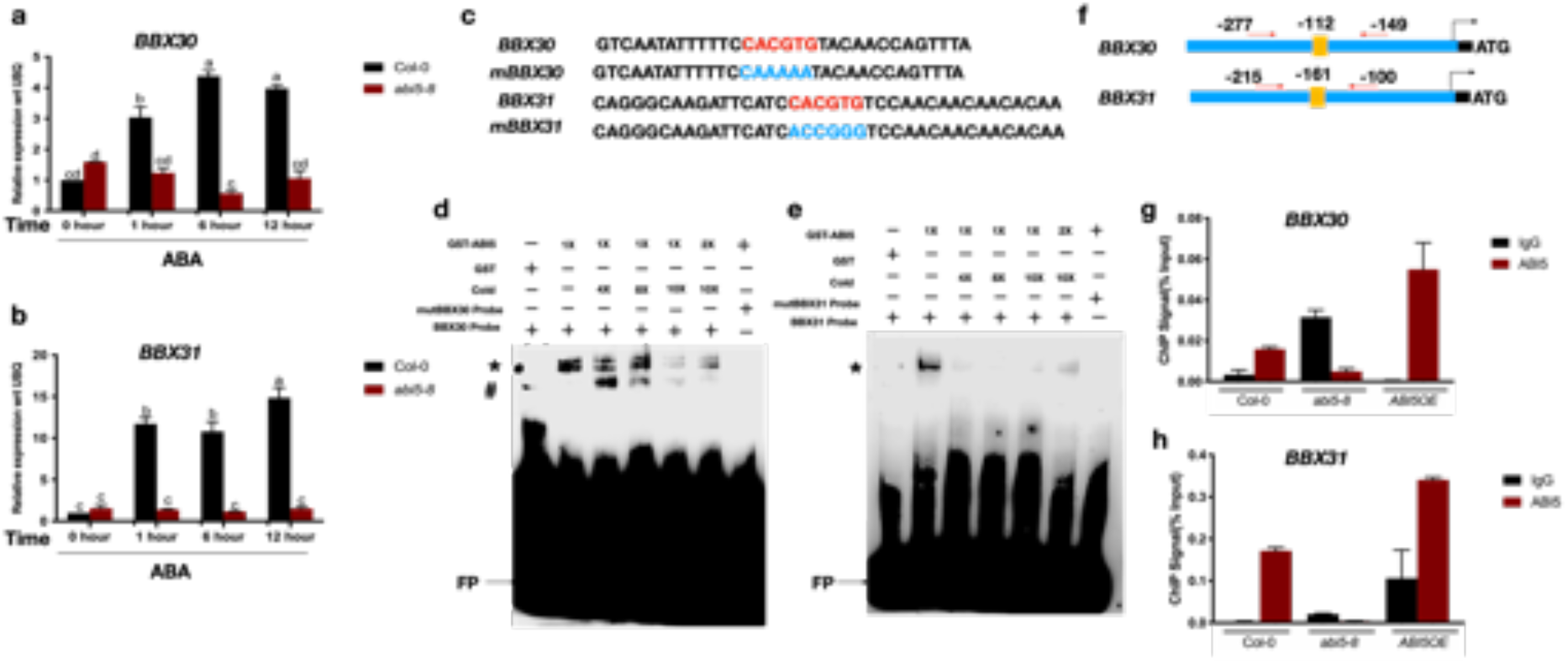
ABI5 directly binds to the promoter of *BBX30* and *BBX31* and induces their transcription. (a-b) The relative expression level of (a) *BBX30* and (b) *BBX31* in the Col-0 and *abi5-8*. Seeds were germinated on 0.5xMS -sucrose conditions. 100% germinated seeds were transferred to medium containing 10μM ABA. Error bars indicate SEM of two biological replicates, the letters above the bars represent different statistical group. (c) Sequence of the probes used for EMSA, mutated nucleotides are shown in blue colour. (d-e) EMSA showing the binding of ABI5 to the promoter of *BBX30* and *BBX31*, containing the G-box domain and labelled with biotin. *mutBBX30* and *mutBBX31* represent the mutated G-box, while + and - indicate presence and absence; cold represents the unlabelled probes used in 4X, 8X, and 10X concentrations. FP represents a free probe. # Denotes the nonspecific shift and * indicates the ABI5 mediated shift. (f) Schematic representation of *BBX30* and *BBX31* promoter. Arrow represents the transcription start site. The yellow boxes indicate G-box type ABRE *cis*- regulatory elements, and the number represents the base location of G-box type elements. (g-h) ChIP-qPCR analysis determining the binding of ABI5 onto the promoter of *BBX30* and *BBX31* in seedlings of Col-0, *abi5-8*, and *ABI5OE*. The genotypes were grown in 1um ABA and samples were harvested on fourth day after stratification. DNA-Protein complexes were immunoprecipitated using anti-ABI5 and anti-IgG antibodies (negative control). ChIP DNA was quantified using primers flanking the G-box region of *BBX30* and *BBX31*. Error bar represents the the mean ± SEM of three technical replicates.

**Fig. 9.**
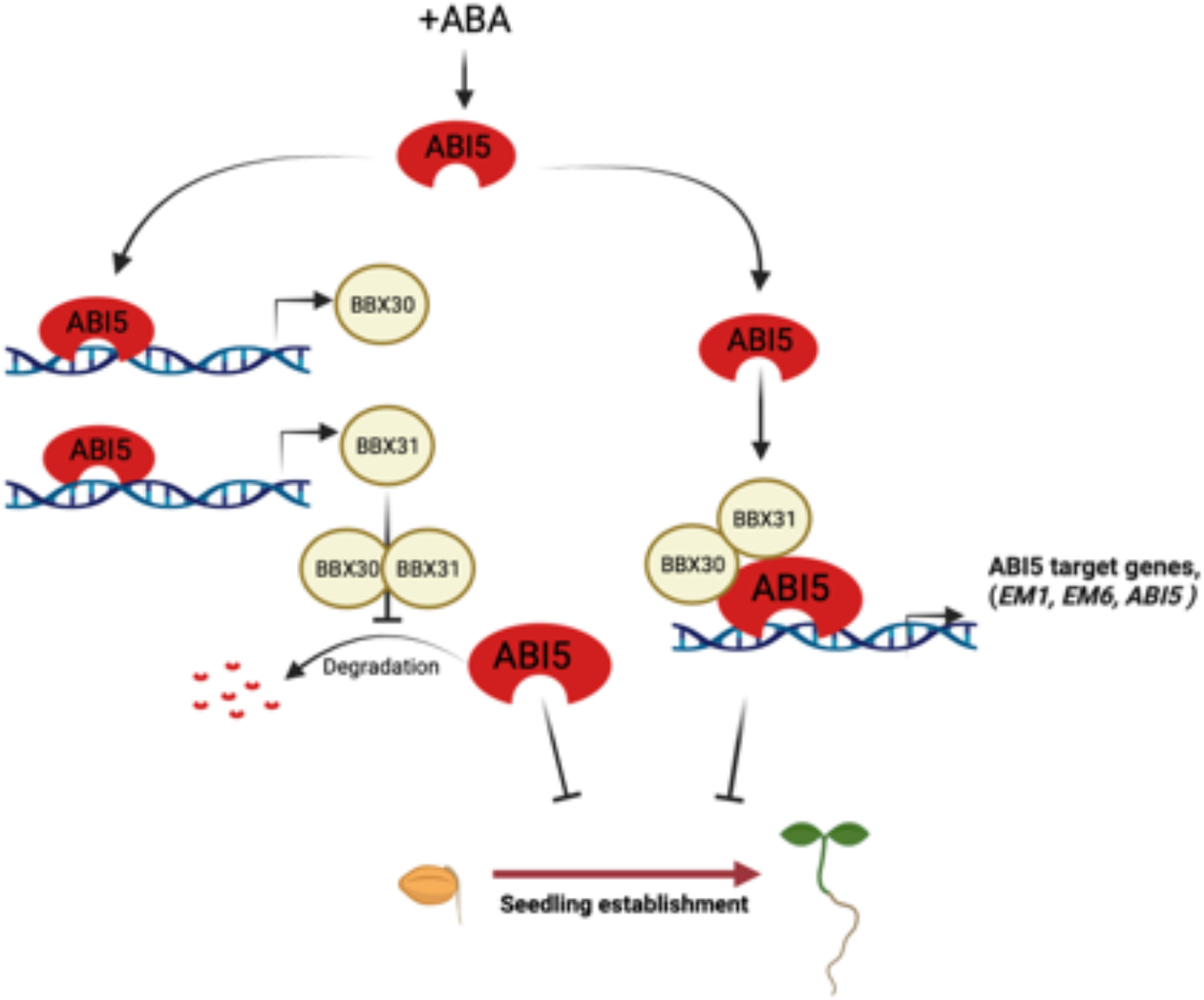
Model showing BX30, BBX31 and ABI5 regulate post-germination seedling development in an interdependent manner. BBX30 and BBX31 promote the stability of ABA-induced ABI5 by inhibiting its degradation and regulating its activity to promote expression of ABI5 target genes. ABI5 on the other hand directly binds to the promoter of *BBX30* and *BBX31* and promotes their transcription. The microproteins miP1a/BBX31 and miP1b/BBX30 alongwith ABI5 thereby form a positive feedback loop to inhibit seedling establishment.

## Discussion

Germination, seedling development and stomatal activity are among the several important developmental events regulated by both ABA and light(Yadukrishnan and Datta, 2021; Ali et al., 2020; Xu et al., 2014; Chen et al., 2008; Lau and Deng., 2007). Of these the integrated role of ABA and light in regulating early seedling development is relatively less known. ABA inhibits post-germination seedling establishment and the ABA sensitivity of seedlings is known to be modulated by light. ABI5 seems to play a key role in mediating the interplay between the light and ABA signaling pathways (Xu et al., 2014; Collin et al., 2021). Several members of the light signaling pathway that are modulated by the light or dark cue, feed the light information either upstream or downstream of ABI5. A number of light signaling factors have been identified to bind to the promoter of *ABI5* and regulate its transcription. These include transcription factors like PIFs, DET1, HY5, BBX19, BBX21, FAR1 and FHY3(Sakuraba et al., 2014; Penfield et al., 2010; Shi et al., 2013; Xu et al., 2014; Kang et al., 2018; Bai et al., 2019). Some other light signaling regulators can modulate ABI5 activity at the protein level. This may be as a result of direct physical binding with ABI5 as in the case of PIF1 or indirectly modulating ABI5 stability by binding to its repressor ABD1 as mediated by COP1(Peng et al., 2022). The stabilization activities of PIF1 and COP1 are limited to the dark environment. COP1 also promotes ABI5 mediated downstream gene activation to suppress post-germination seedling establishment (Yadukrishnan et al., 2020). PIF1 and ABI5 act in a cooperative manner to activate common downstream target genes by binding to their promoters in the dark(Kim et al., 2016).To prevent precocious seedling establishment ABI5 needs to be active both under dark and in light. Here we identified two microProteins miP1a/ BBX31 and miP1b/BBX30 that directly interact with ABI5 to stabilize it and promote its downstream activities in the light (Fig. 5, 6, 7). ABI5 via a positive feedback loop regulates the transcription of miP1a/BBX31 and miP1b/BBX30 (Fig. 8). The microProteins and ABI5 thus mediate seedling growth arrest in an interdependent manner, at least partially (Fig. 1, 2, 3, 4). This study provides a novel microprotein mediated post-translational regulatory mechanism for controlling ABI5 accumulation and activity during post-germination seedling establishment in the absence of COP1and PIFs that are deactivated in light.

BBX proteins are zinc-finger transcription factors that regulate numerous aspects of early seedling development (Yadav et al., 2020, 2020; Song et al., 2020). In *Arabidopsis* there are 32 B-box proteins that are divided into five structural groups based on the domain structure(Gangappa and Botto, 2014). The structural groups I-III (BBX1-BBX17) contain a CCT domain at the C-terminal half which is often involved in DNA binding and transcriptional regulation. BBX11 regulates the accumulation of protochlorophyllide to optimize greening during de-etiolation(Job, 2020). Structural group IV contains 2 B-box domains but is devoid of the CCT domain. Four members of this group (BBX20, BBX21, BBX22, BBX23) have been characterized as positive regulators of photomorphogenesis while the other four (BBX18, BBX19, BBX24, BBX25) repress photomorphogenic development. BBX20, BBX21, BBX22 act as cofactors of HY5 to mediate light-regulated development (Bursch et al., 2020; Job et al., 2018). An M6 motif identified in the C-terminus of these proteins might regulate their specific developmental responses in response to light (Yadukrishnan et al., 2018). Structural group five consists of 7 members (BBX26-BBX32) that contain only one B-box domain. Functional characterization of these single-domain BBX proteins indicates that many of them play important roles in early seedling development. BBX30 and BBX31 promote hypocotyl elongation in seedlings under visible light(Yadav et al., 2019; Heng et al., 2019). HY5 inhibits the transcription of *BBX30* and *BBX31* (Heng et al., 2019). Interestingly, BBX30 and BBX31 positively regulate *BBX28* and *BBX29* which suppresses HY5-mediated inhibition to promote the transcription of *BBX30* and *BBX31* to fine-tune photomorphogenic development (Song et al., 2020). BBX32 interacts with BBX21 and suppresses the promotion of light-mediated gene expression by BBX21 and HY5(Holtan et al., 2011). Wu et al elegantly characterized the specific role of BBX30 and BBX31 during de-etiolation (Wu et al., 2020). These two proteins inhibit the dimerization of PIF3 and EIN3 to promote apical hook and cotyledon opening during the dark to light transition. It is interesting to note that other group V members BBX32, BBX28 and BBX29 promote BR-mediated cotyledon closure in the dark (Ravindran et al., 2021; Cao et al., 2022). The functional diversity between these closely related BBX proteins needs further characterization. Our study here shows that BBX30 and BBX31 inhibit post-germination seedling establishment in an ABA-dependent manner. It seems that the same proteins may be recruited for different functions depending on the presence or absence of stress conditions. In addition to their role in modulating early seedling development, BBX30 and BBX31 also regulate flowering and BBX31 promotes UV-B stress tolerance(Yadav et al., 2019; Graeff et al., 2016).

MicroProteins miP1a/BBX31 and miP1b/BBX30 are 121 and 117 amino acids long respectively and show 65.5% sequence identity(Graeff et al., 2016). microProteins form homodimers, heterodimers, or multimeric complexes by interacting with the protein-protein interaction domains of their targets. As of now, not more than 50 miPs have been functionally characterized in plants. In most of these cases, the microProteins repress the activity of their targets(Staudt and Wenkel, 2011). In fact, miP1a and miP1b have been previously shown to form a repressor complex together with CO and TOPLESS to inhibit flowering(Graeff et al., 2016). This microProtein duo also inhibits the oligomerization of PIF3 and EIN3 to repress their transcriptional activation potential (Wu et al., 2020). Here we report that miP1a/BBX31 and miP1b/BBX30 physically interact with ABI5, stabilize it and promote its binding to the downstream targets to activate their expression (Fig. 5,6,7). The versatility in the mode of action of the same miPs probably depending on the developmental stage of the plant and the environmental conditions is intriguing. During dark to light transition, miP1a/BBX31 and miP1b/BBX30 promote seedling de-etiolation, however, in the presence of ABA, these miPs arrest seedling development. What is the basis of this differential regulation in the presence or absence of stress is yet to be discovered. Our data shows that ABA induces the expression of *miP1a/BBX31* and *miP1b/BBX30* in a ABI5 dependent manner (Fig. 8). ABI5 binds to the G-box motif present in the promoters of *miP1a/BBX31* and *miP1b/BBX30* to activate their expression (Fig. 8). Previously HY5 has been shown to bind to the G-box of *BBX30* and *BBX31* to repress their transcription in light(Heng et al., 2019; Yadav et al., 2019). However this differential transcriptional regulation of the miPs is unlikely to account for their varied post-translational activity in presence or absence of ABA. The upregulation of *miP1a/BBX31* and *miP1b/BBX30* expression is extremely rapid and transient happening within the 1 hour of the transfer from dark to light (Wu et al., 2020). The inhibitory activity of miP1a/BBX31 and miP1b/BBX30 in preventing PIF3 and EIN3 functional oligomer formation also seems to be transient as PIF3 and EIN3 are eventually degraded by light. It is possible that the miPs might disengage from this transient activity in the presence of stress. It would be interesting to see what happens to the allosteric deactivation of PIF3 and EIN3 by miP1a and miP1b in presence of ABA. The cooperative action of miP1a and miP1b in interacting with its targets is also a unique feature of this regulatory module that needs further characterization. In several eukaryotes, miPs regulate transitions in the cell cycle and circadian clock. In yeast, the microProtein Nrs1 rewires the transcriptional machinery to regulate G1/S transition under nutrient stress conditions. This study and some previous reports indicate the role of miP1a/BBX31 and miP1b/BBX30 in modulating the transition from seed to seedling and vegetative to flowering state (Wu et al., 2020; Graeff et al., 2016). In the plant kingdom, miPs are generally present in dicotyledonous flowering plants and might have evolved to modulate these crucial transitions that regulate plant life and productivity (Graeff et al., 2016). The identification of light regulated microProteins in modulating ABA sensitivity during the early establishment phase, might open the doors for optogenetic manipulation of these versatile regulators. In the future, synthetic miPs may be used as molecular brakes to postpone development during unfavourable conditions to ensure plant survival.

## Acknowledgements

DS acknowledges IISER Bhopal for PhD fellowship. SD acknowledges the SERB grant STR/2021/000046 for funding. Authors acknowledge Dr. Stephan Wenkel and Dr. Arpita Yadav for providing *miP1b*/*BBX30* and *miP1a*/*BBX31* mutant and overexpressor lines used in previous studies. Authors also acknowledge Dr. Nevedha Ravindran, Rahul Puthan Valappil and Dr. Yadukrishnan Prem for sharing *ABI5* constructs and *ABI5OE* lines. Authors acknowledge the assistance provided by Rucha Kulkarni and Akshat Singh Raghuwanshi for some expreiments. Authors thank Henrik Johansson and Yadukrishnan Prem for critical comments. Authors thanks all current PCDB lab members especially Dr. Amit Kushwaha, Arpan Mukherjee, Shubhi Dwivedi, Debojyoti Kar, Rahul Puthan Valapil and Ajar Anupam Pradhan for critical reading of the manuscript and providing valuable comments.

## Author contributaion

SD and DS conceived the project. DS planned and performed all the experiments. SD wrote the manuscript with help from DS. DS and SD analysed the data and helped preparing the final manuscript.

## Data Availability Statement

The data that supports the findings of this study are available in the supplementary material of this article.

