## supplemental figures 1-6, table 1 for "MicroProteins miP1b/BBX30 and miP1a/BBX31 form a positive feedback loop with ABI5 to retard seedling establishment"

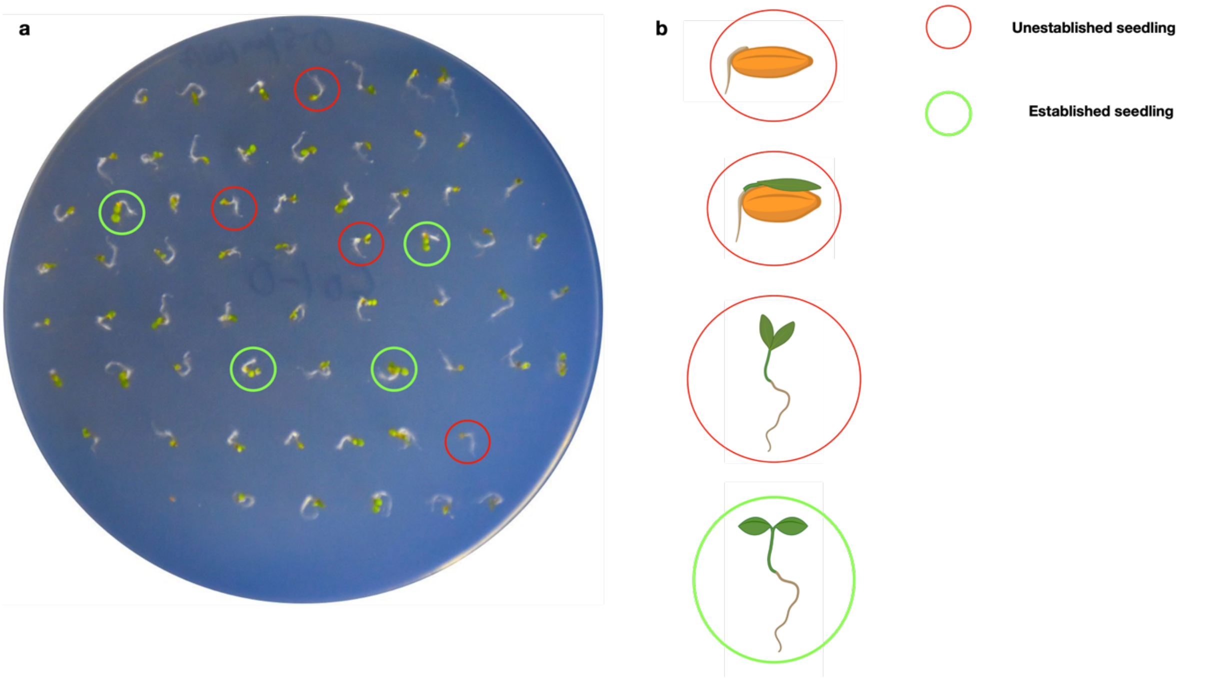

**Fig. S1 Arabidopsis seedlings with open, expanded green cotyledons were counted as established seedlings.** (b) Plate showing 6-d old Col-0 seedlings growing on 0.5 µM ABA. Green encircled seedlings with open, expanded green cotyledons are established seedlings while red encircled seedlings are not counted as established. (b) Schematic diagram showing the seedling development from germination to complete establishment.

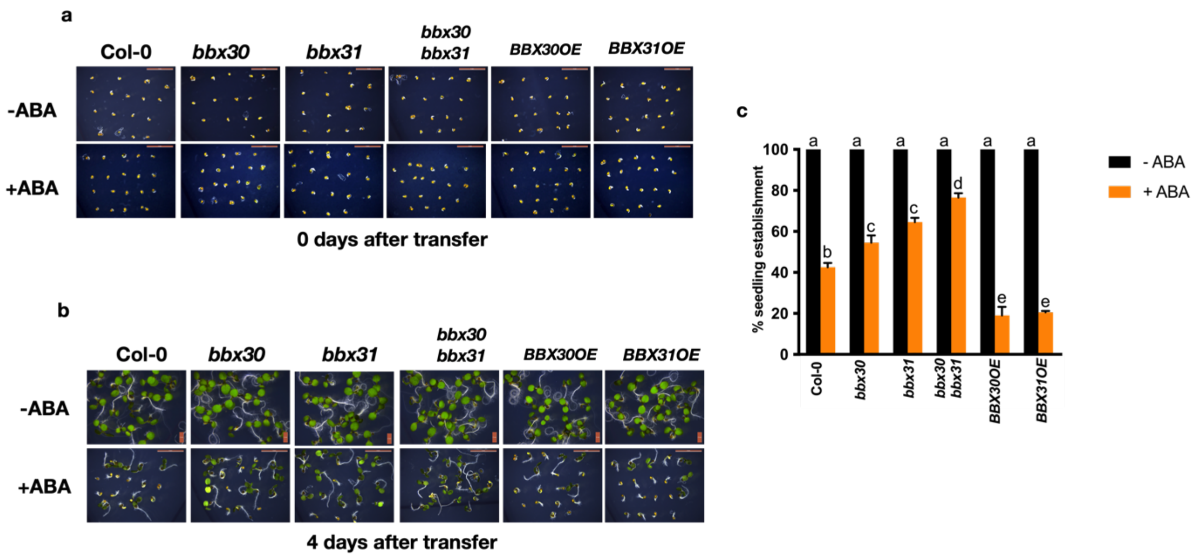

**Fig. S2 Seedling establishment of the indicated genotypes when ABA was provided after germination.** (a-b) Representative images of Col-0, *bbx30, bbx32, bbx30bbx31, BBX30OE, BBX31OE* germinated on 1/2MS plate and transferred to 1/2MS plate supplemented with or without ABA, (a) Imaged just after transfer (b) Imaged 4 days after transfer. (c) Seedling establishment rate of indicated genotypes 4 days after their transfer to –ABA and +ABA plates.

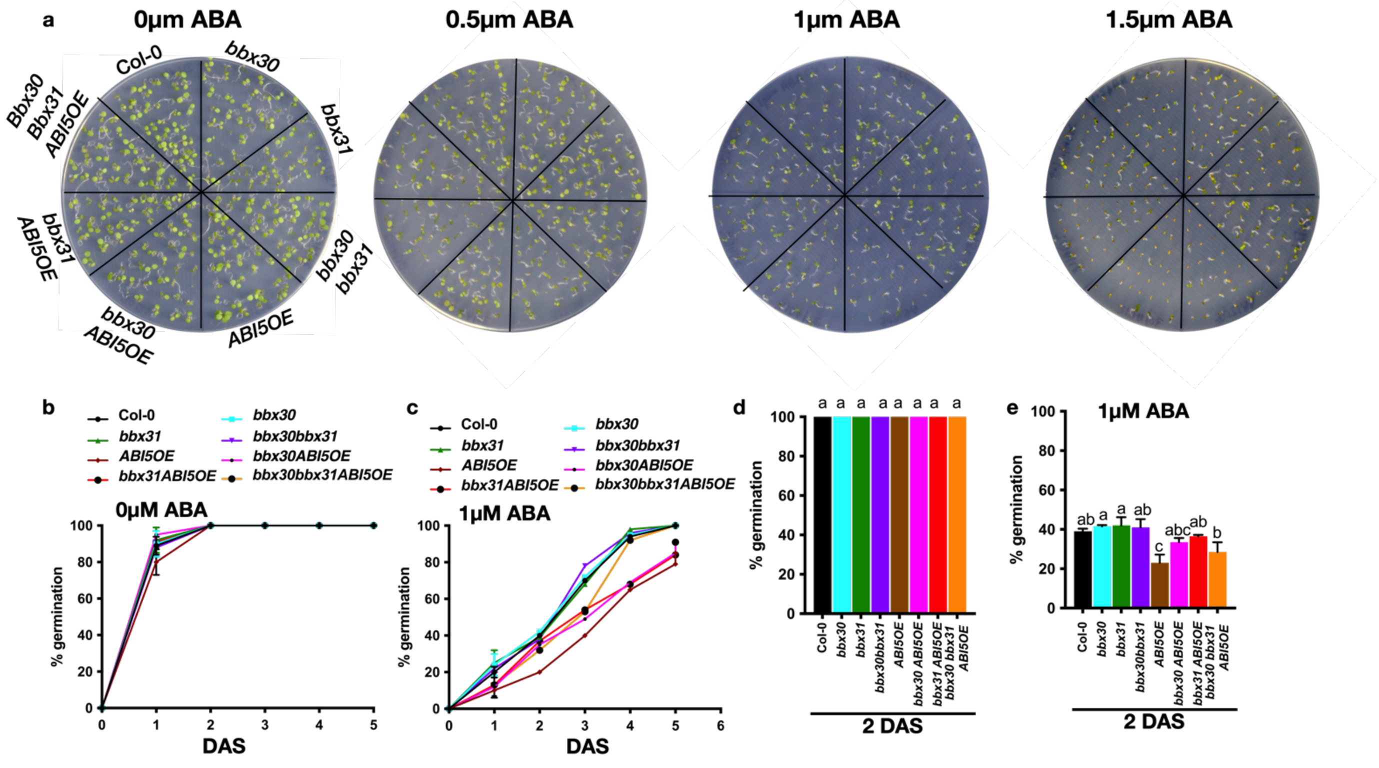

**Fig. S3 % Germination of Col-0*, bbx30 bbx31, bbx30, bbx31*, *ABI5OE*, *bbx30 ABI5OE, bbx31 ABI5OE*, *bbx30 bbx31 ABI5OE* in +/- ABA conditions**

(a) Plate images of 6-day-old seedlings of Col-0*, bbx30, bbx31*, *bbx30bbx31, ABI5OE*, *bbx30ABI5OE, bbx31ABI5OE*, *bbx30bbx31ABI5OE* grown on a 0.5x MS plate supplemented with 0µM, 0.5µM, 1µM, 1.5µM of ABA (b-e) % germination in the indicated genotypes under (b,d) 0µM and (c,e) 1µM ABA conditions. DAS indicates days after stratification. In (d,e) error bar represents SEM of three independent experiments with >500 seeds. Letters above the bar indicate the statistical groups as determined by one-way ANOVA followed by Tukey's *post hoc* test ($p < 0.05$).

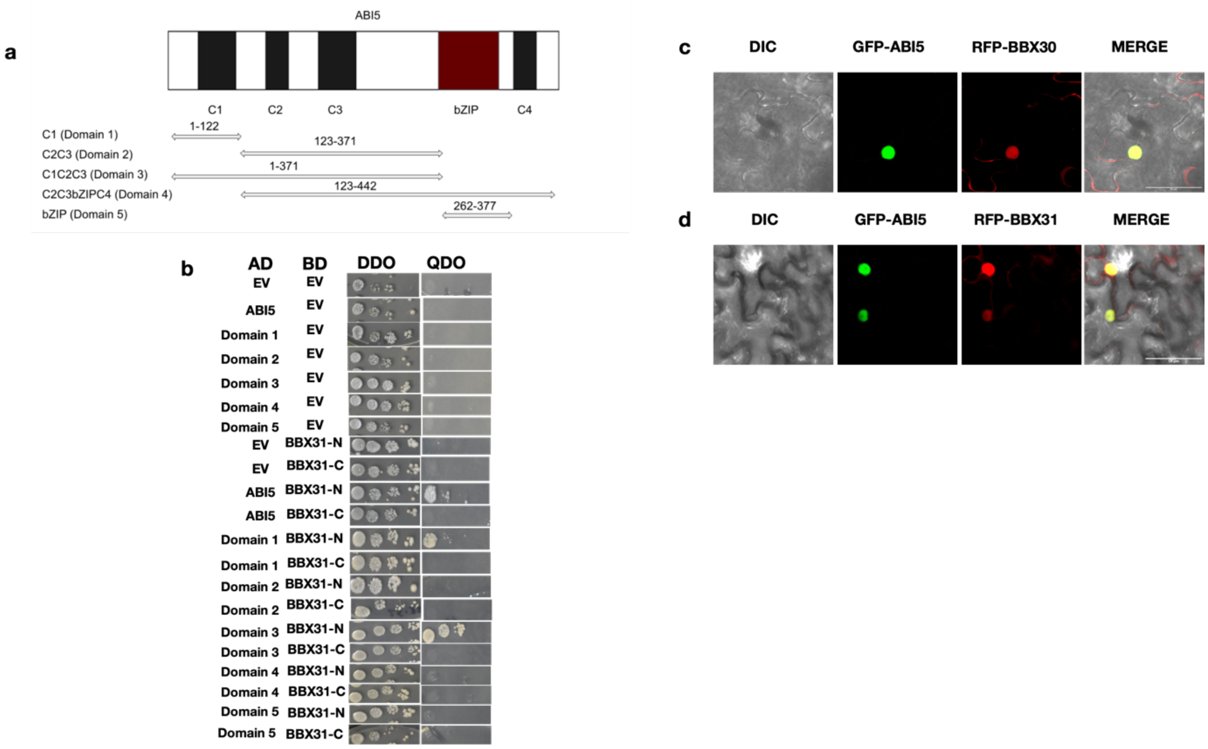

**Fig. S4 Physical interaction between BBX31 and ABI5.**

(a-b) Yeast two-hybrid assay showing the interaction between the N-terminal and C-terminal domain of BBX31, full length ABI5 and different domains of ABI5 as shown in (a). AD and BD represent the GAL4 activation domain and binding domain. N terminal and C-terminal half of BBX31 are fused to AD, while ABI5 and its domains were fused with BD domains and tested for interaction. DDO represents the medium lacking leucine and tryptophan, while QDO additionally lacks histidine and adenine. (c-d) Colocalization of BBX30 and BBX31 with ABI5 in the nucleus of *N. benthamiana* leaf epidermal cells. BBX30 and BBX31 were fused with RFP and ABI5 is fused with GFP, Both the constructs were transformed into *Agrobacterium* and co-infiltrated into 3-week-old leaves and images were captured after 2 d. The scale bar measures 50 µm

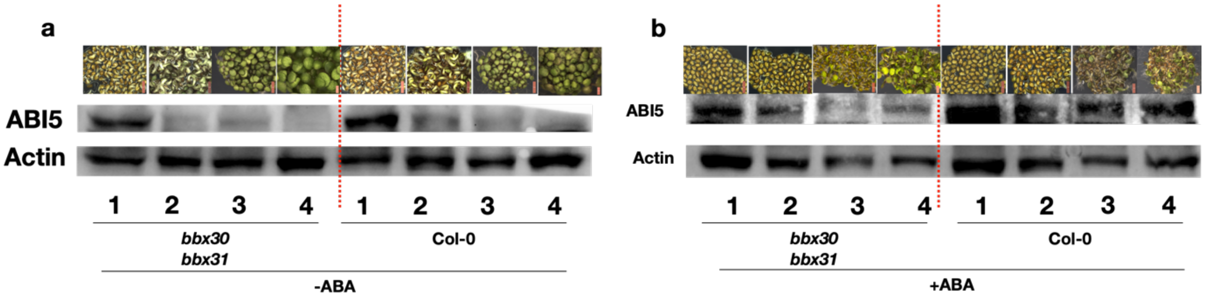

**Fig. S5 BBX30 and BBX31 promote ABI5 accumulation upon ABA treatment**
(a-b) Immunoblot showing ABI5 accumulation in *bbx30 bbx31* and Col-0 in (a) -ABA and (b) +ABA conditions. Top row shows image of seeds/seedlings inoculated on 0.5x MS-sucrose plates with or without 1µM ABA. Samples were harvested at day 1, 2, 3, and 4 followed by immunoblotting using anti-ABI5 antibody. Actin was used as an internal control.

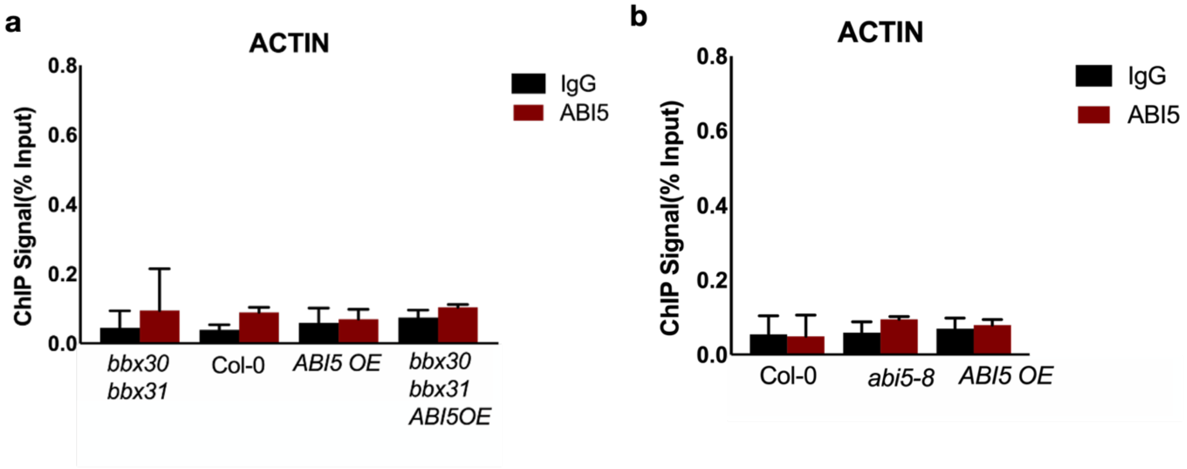

**Fig. S6 ChIP-qPCR showing the lack of ABI5 enrichment over *ACTIN* promoter**
(a-b) Experimental conditions were same as Figure 7, I and J and Fig. 8 g and h. ChIP DNA was quantified using *ACTIN* promoter specific primers.

### Table S1:

| | |
|---|---|
| Primers for gateway cloning | |
| BBX30 B1 | GGGGACAAGTTTGTACAAAAAAGCAGGCTTCATGTGTAGAGGGTTTGAGAAAGAAG |
| BBX30 B2 | GGGGACCACTTTGTACAAGAAAGCTGGGTCGAGAAACACAAAGGGAATTTTGTG |
| BBX31 B1 | GGGGACAAGTTTGTACAAAAAAGCAGGCTTAATGTGTAGAGGCTTGAATAATGA |
| BBX31 B2 | GGGGACCACTTTGTACAAGAAAGCTGGGTCTCAGAGAAAAACAAACGGAACC |
| BBX31 NterminalB2 | GGGGACCACTTTGTACAAGAAAGCTGGGTTGCGCCTTACGTGTCTCCAAGCT |
| BBX31 CterminalB1 | GGGGACAAGTTTGTACAAAAAAGCAGGCTTAGTGCTATGCACTTCTTGTCAGA |
| ABI5_Gw_F | GGGGACAAGTTTGTACAAAAAAGCAGGCTTCATGGTAACTAGAGAAACGAAGT |
| ABI5_Gw_R | GGGGACCACTTTGTACAAGAAAGCTGGGTCCTTAGAGTGGACAACTCGGG |
| Primers for genetarting protein constructs | |
| ABI5 FW | CGCGAATTCATGGTAACTAGAGAAACGAAGTTG |
| ABI5 RW | CGCCTCGAGTACAAGAAAGCTGGGTGTTAGAG |
| BBX30 FW | CGCGGATCCATGTGTAGAGGGTTTGAGAAAG |
| BBX30 RW | CGCCTCGAGTCAGAGAAACACAAAGGGAATTTTG |
| BBX31 FW | CGCGGATCCATGTGTAGAGGCTTGAATAATGAAG |
| BBX31 RW | CGCCTCGAGTCAGAGAAAAACAAACGGAACC |
| Primers for EMSA | |
| bbx31FW | AAATAACAGGGCAAGATTCATCCACGTGTCCAACAACAACACAACACAAA |
| bbx31 RW | TTTGTGTTGTGTTGTTGTTGGACACGTGGATGAATCTTGCCCTGTTATTT |
| bbx31 mutFw | AAATAACAGGGCAAGATTCATCACCGGGTCCAACAACAACACAACACAAA |
| bbx31 mut Rw | TTTGTGTTGTGTTGTTGTTGGACCCGGTGATGAATCTTGCCCTGTTATTT |
| BBX30 Fw | GTCAATATTTTTCCACGTGTACAACCAGTTTA |
| BBX30 Rw | TAAACTGGTTGTACACGTGGAAAAATATTGAC |
| BBX30 mutFw | GTCAATATTTTTCCAAAAATACAACCAGTTTA |
| BBX30 mutRw | TAAACTGGTTGTATTTTTGGAAAAATATTGAC |
| EM6 FW | AATAAAGTCAGACACGTGGCATGTCACCAA |
| EM6 RW | TTGGTGACATGCCACGTGTCTGACTTTATT |
| ABI5 FW | GCAAGAAACAGAAGAGAGACACGTGCAGGACACGTGTCGTCCGCAGCCGAACGGA |
| ABI5 RW | TCCGTTCGGCTGCGGACGACACGTGTCCTGCACGTGTCTCTCTTCTGTTTCTTGC |
| Primers for ChIP | |
| BBX30 Fw | GGAGGGAAAATAACTAAATTATTG |
| BBX30 Rw | CTTGTGTTCAAGATTTAGGTTTA |
| BBX31 Fw | TGTGCGTCCAATATGAGGTC |
| BBX31 Rw | TGATGTGTTTGTGTTGTGTTGTT |
| ACTIN 8 Fw | GCCTCTGTAAATCAAAACCCCA |
| ACTIN 8 Rv | TCTTTTCGCAGGAACCCAATT |
| EM1 ChIP Fw | GGATTAAGATTAATCGGAGTCG |
| EM1 ChIP Rw | GTGGAAGAGAAGACGCGGCGAG |
| EM6 ChIP Fw | GCGGCGGTATAGTTAAAGAACA |
| EM6 ChIP Rw | GATGATATACGAAGAAGACT |
| Primers for genotyping | |
| LBb1.3 | ATTTTGCCGATTTCGGAAC |
| abi5-8-_LP | CAATGGAAGTTCGGAATCATG |
| abi5-8 RP | CACTCGTTTTCTTCTTAAAGCG |
| BBX31- CRISPR Fw | GCAGAAGAAGTGACGGAGGA |
| BBX31- CRISPR Rv | AGAAAAACAAACGGAACCTCA |
| BBX31- CRISPR FW | GAAGAAGCGACAATGGAGGATG |
| BBX31- CRISPR RW | ACGAGTTAGCTTCCGACAGG |
| Primers for qPCR | |
| ABI5_QF | GAGAATGCGCAGCTAAAACA |
| ABI5_QR | GTGGACAACTCGGGTTCCTC |
| EM1_QF | CGAGCTACTAGTGTCCGCTGCA |
| EM1_QR | GTAAAACCAACCGGCAACCGCA |
| EM6_QF | CTTGTCTCCGGTGCTAAG |
| EM6_QR | CAACAGCATCTCGCTGAAG |
| UBQ10_QF | GGCCTTGTATAATCCCTGATGAATAAG |
| UBQ10_QR | AAAGAGATAACAGGAACGGAAACATAGT |
| BBX30_QF | ATGTGTAGAGGGTTTGAGAAAGA |
| BBX30_QR | TGCGTCTGCCTCACAATACA |
| BBX31_QF | ATGTGTAGAGGCTTGAATAATGAAGAG |
| BBX31_QR | TCACATTTTCTACAGAGGAACGC |
